# Circadian Modulation of Neutrophil Function Determines Collateral Perfusion and Outcome After Ischemic Stroke

**DOI:** 10.1101/2025.03.09.639714

**Authors:** Sandra Vázquez-Reyes, Alicia García-Culebras, Gaohong Di, Blanca Díaz- Benito, Francisco J. De Castro-Millán, Alessandra Ruiz-Sanchez, Eneko Merino-Casamayor, Carlos Parra-Pérez, Carmen Nieto-Vaquero, Ana Moraga, Patricia Calleja, Ana Dopazo, Sergio Callejas, Andrea Rubio-Ponce, Alejandra Aroca-Crevillén, Fátima Sánchez-Cabo, Sara Pascual El Bobakry, Carlos Torroja, Elga Esposito, Eng H. Lo, Iván Ballesteros, Andrés Hidalgo, María Isabel Cuartero, Ignacio Lizasoain, María Ángeles Moro

**Author notes:** Equal contribution (Sandra Vázquez-Reyes, Alicia García-Culebras, Gaohong Di, Blanca Díaz- Benito). Correspondence to: María A. Moro, Maria I. Cuartero or Ignacio Lizasoain., G. Di’s present address: Anesthesiology Department, Sixth Hospital Wuhan, Jianghan University, China.

## Abstract

Stroke is a leading cause of mortality and disability, driven by complex and time-dependent mechanisms that aggravate ischemic damage. Among them, collateral perfusion determines the initial size of the ischemic core, the rate of its expansion, and the extent of the penumbra both at stroke onset and over time. Insufficiency of collaterals may occur due to genetic factors or other determinants, such as aging or cardiovascular risk factors, which reduce the number of collaterals or the diameter of those that remain. But aspects of less structural nature could also affect the effectiveness of these pathways by decreasing their patency. We hereby show that diurnal fluctuations in infarct volume in ischemic stroke mouse models are neutrophil phenotype-dependent, since differences in infarct volumes were abolished by depleting neutrophils or blocking their circadian clock, and linked to the collateral circulation: during the inactive phase of mice (daytime), collateral perfusion in the ipsilesional hemisphere was reduced, coinciding with an increase in intravascular neutrophil accumulation, suggestive of microvascular stalling. Single-cell transcriptomics, *ex vivo* functional assays and *in vivo* pharmacological and genetic strategies confirmed enhanced neutrophil extracellular traps (NETs) formation at this time. Importantly, in a cohort of human stroke patients, we identified diurnal oscillations in neutrophil and NET-related biomarkers, peaking during the human inactive phase (evening/night), and similarly associated with reduced collateral flow and poorer clinical outcomes. These findings underscore the critical role of neutrophils, their circadian dynamics and NET release in driving collateral insufficiency and ischemic brain damage, suggesting novel personalized therapeutic strategies based on circadian rhythms for the treatment of stroke.

## INTRODUCTION

Ischemic stroke is a devastating disease with high morbidity and mortality rates, which highlights the need for new therapies to improve patient outcomes. Despite extensive exploration of neuroprotective treatments for acute stroke, none have demonstrated sufficient efficacy to be translated into clinical practice. While these treatments often seem very promising in animal models, they consistently fall short in clinical settings, highlighting a critical gap in stroke therapy^1^. Circadian rhythms, with a crucial role in the timing and outcomes of ischemic stroke^2,3^, have been identified as a key factor that could explain the translational failure of these pharmacological strategies^4^. Circadian rhythms can influence the progression and outcome of stroke by influencing crucial aspects of the ischemic cascade and post-stroke damage, such as the inflammatory response^5–9^. The immune system is highly circadian-regulated with a 24-hour rhythmicity both at steady state and during activation. Neutrophils, the first immune responders in post-stroke inflammation, exhibit circadian fluctuations in number, phenotype, and activation markers potentially influencing stroke outcomes based on time of day^10,11^. Neutrophils exert damage after stroke by different mechanisms, including oxidative stress or blood-brain barrier (BBB) disruption. Importantly, they are also key contributors to the development of the no-reflow phenomenon and immunothrombosis through the release of neutrophil extracellular traps (NETs)^12–14^. NETs are networks of decondensed DNA combined with histones and neutrophil proteases, such as elastase (NE) and myeloperoxidase (MPO), released by neutrophils during immune activation^14,15^. Although the main functions of NETs include capturing and killing pathogens them through the action of antimicrobial peptides, they can also contribute to inflammation and tissue damage in sterile conditions, including stroke, by interacting with platelets and promoting microthrombosis, or by directly contributing to neuronal death^16,17^.

Based on this, neutrophils are key mediators of damage following a stroke. Congruently, many studies have shown that neutrophil depletion and/or inhibition of their infiltration may improve stroke outcome after experimental stroke in mice^18,19^; however, this neuroprotective effect is not always consistent, reflecting that, as demonstrated in both physiology and different pathological contexts^20^, neutrophils are a heterogeneous population that may acquire different phenotypes after stroke by mediating both damage and tissue repair depending on their phenotype^21–24^. A source of this functional heterogeneity after cerebral ischemia may arise from circadian rhythms. Indeed, previous studies have described that neutrophils undergo diurnal aging in the circulation, resulting in changes in gene expression and functional properties, such as ROS production, migration and capacity to produce NETs^6,10,11^.

This study aimed to gain insight into the neutrophil-dependent mechanisms that mediate damage after stroke and to explore its circadian influence using a murine model of thrombotic stroke and in stroke patients. Our data demonstrate that neutrophils may adopt different phenotypes after stroke and, importantly, that this functional heterogeneity is regulated by circadian rhythms and account for stroke outcome both in mice and humans. Our results also show that stroke-associated NET release is a key property influenced by circadian rhythms that drives stroke pathophysiology. We propose that neutrophil-mediated capillary stalling under the influence of circadian rhythms is a relevant target for precise and efficient therapeutic approaches against stroke.

## METHODS

### Animals

All experimental protocols adhered to the guidelines of the Animal Welfare Committee of the CNIC and the Ministry of Environment and Territorial Planning of the Community of Madrid (RD 53/2013; PROEX 047/16), in compliance with European directives 86/609/EEC and 2010/63/EU. Furthermore, the procedures are reported following the ARRIVE (Animal Research: Reporting of In Vivo Experiments) guidelines^25^. The experimental groups were housed in the CNIC animal facility. The animals were kept under standard temperature and humidity conditions, with a 12-hour light-dark cycle. Food and water were provided ad libitum, both inside and outside the inverted light cabinet. Experiments were performed in 8-to 12-week-old male C57BL/6 mice and 12 months, in the case of the aged group. The following knock-in or transgenic mice were used: Bmal^fl/fl^MRP8^Cre^ or Bmal1^Neu^ (mice with neutrophil-specific deficiency in Bmal1)^26,27^, Ly6G^tdTom^ (mice with high levels of tdTomato fluorescence in neutrophils)^28^ and Cxcr4^fl/fl^MRP8^Cre^ or Cxcr4^Neu^ (mice with neutrophil-specific deficiency in CXCR4). We also used PAD4^−/−^ (mice with lack exons 9-10 of the peptidyl arginine deiminase type IV gene) obtained from The Jackson Laboratory (Bar Harbor, Me), and Dnase1^−/–^ Dnase1l3^−/–^ or D1^−/−^D1l3^−/−^, (mice without anti-dsDNA reactivity) kindly provided by Markus Napirei^29^. All transgenic mice were in the C57BL/6 background.

### Experimental models of cerebral ischemia

Animals were randomly assigned to experimental groups using a randomized fashion (coin toss). Mice were anesthetized with sevoflurane 2-3% in a mixture of 80% air/20% oxygen; body temperature was maintained at physiological levels with a heating pad during the process. In surgical procedures a scalp incision was made, and the right temporal muscle was dissected. The area between zygomatic arch and squamous bone was thinned by a high-speed drill and cooled with saline. MCA trace was visualized with a stereomicroscope (PZMIV, World Precision Instruments, USA) and a thin bone film over MCA was lifted up.

In the *ligature model by permanent or transient middle cerebral artery occlusion* (pMCAO or tMCAO), left MCA was occluded by ligature of the trunk just before its bifurcation, between the frontal and parietal branches, with a 9-0 suture (61966, Lorca Marin) after removing the meninges in that area^30,31^. In the transient occlusion, MCA were reperfused after 75 minutes occlusion by removing the knot^32^.

The *ferric chloride (FeCl_3_) permanent stroke model* was performed as described^33^. A piece of Whatman filter paper strip soaked in freshly prepared ferric chloride (20%) was placed over the intact duramater in the region of the MCA for 10 minutes and then removed to allow the formation of a thrombus.

The *thromboembolic in situ model* was performed as described^34,35^. This model involves injecting thrombin into the MCA to create a fibrin-rich thrombus that occludes the artery. Thrombin is injected using micropipettes made from graduated capillaries, shaped to prevent bleeding during removal. The micropipette is loaded with thrombin through a pneumatic system and stored at –80°C until use. The MCA is located through a surgical procedure that involves incisions to expose the temporal bone and using a microscope to identify the artery. The micropipette is inserted into the artery lumen to inject 1 μL of thrombin, causing occlusion and a Doppler laser probe is used to measure blood flow. The occlusion is considered stable if there is a significant and sustained reduction in cerebral blood flow, measured by Doppler, and changes in vessel color. Partial or complete flow recovery in any branch is considered reperfusion.

In all types of surgeries, mice in which the MCA was exposed but not occluded served as sham-operated controls. In several experiments, naïve animals that did not undergo any type of surgery were used as control.

### Infarct size determination

Infarct volume was assessed at 24h using magnetic resonance imaging (MRI) system (7T-MRI Agilent-Varian), by Nissl staining and by TTC staining (Sigma-Aldrich, T8877-50G). From Nissl staining, the slides were first digitized using the AxioScan Z1” (Zeiss®) at 40x magnification. Subsequently, 14 rostro-caudally ordered slices were measured using the NDP.view 2.7.39+ software (Hamamatsu Photonics K.K.). Infarct size in all cases was determined as described^31^ and was expressed as % of injured ipsilateral hemisphere by the formula (CH-NLH/CH) X 100. The value resulting from this formula is then normalized by the edema index which is the ratio between the volume of the contralateral and ipsilateral hemisphere.

### Extraction and processing of organs for flow cytometry studies

Blood was taken through cardiac puncture with 1ml syringe containing EDTA (0.5M EDTA at 0.1%) attached to a 26g needle filled with 50mL of 0.5M EDTA. Samples were lysed in 1X RBC lysis buffer (diluted from a 10X stock: KH_4_Cl 1.5M, KHCO_3_ 0.1M, EDTA-Na_2_ 1mM adjusted to pH 7.4) for 10min. For BM cells, mice femurs were flushed using a 23G needle in HBSS1X, then disaggregated with an 18G blunt needle and passed through a 70-mm nylon mesh sieve. Lungs were harvested and cut dry into small pieces before digested in liberase TM (Sigma-Aldrich, 05401119001) and DNase-1 (Sigma) for 20 min at 37°C. For the calvaria, after removing the dura mater, the bone was minced with scissors in 2mL of HBSS1X and then filtered through a 70-μm filter. Meninges were removed from the skull with forceps and minced over a 70-μm filter. The infarcted area was dissected and digested with an enzyme cocktail (containing 50 U/ml collagenase, 8.5 U/ml dispase, 100 µg/ml Nα-tosyl-L-lysine chloromethyl ketone hydrochloride, and 5 U/ml DNase I in 9.64 mL HBSS1X (Gibco™ HBSS, no calcium, no magnesium, no phenol red, Fisher Scientific, 14175-095) for 30 min at 37°C. After digestion, the tissue was ground in a 2mL glass-glass grinder, filtered through a 70-μm filter, centrifuged, and the supernatant was discarded. The pellet was resuspended in 7mL of 35% Percoll, with 5mL of HBSS1X carefully layered on top to create a density gradient, and centrifuged at 4°C, 800g, for 30min without braking. After centrifugation, the myelin at the gradient interface, along with the rest of the supernatant, was discarded, leaving a pellet that was resuspended in 100μL of FACS buffer for staining. All the resulting suspension of cells was centrifuged at 1800 rpm, 4°C, for 5 min, the supernatant was discarded, and the pellet was resuspended in FACS buffer.

### Flow cytometry

All cell pellets were resuspended in FACS buffer (DPBS containing 0.1% low endotoxin BSA). Cell suspensions were stained with the following fluorochrome-conjugated monoclonal antibodies for 15 min at a concentration of 1:1000: anti-mouse Ly6G-PE (clone 1A8, BD Bioscience); anti-mouse CD45-FITC (clone 30-F11, BD Bioscience); anti-mouse/human CD11b-Brilliant Violet 510™ (clone M1/70, BioLegend); anti-mouse CD182 (CXCR2)-PerCP/Cy5.5 (clone SA044G4, BioLegend); anti-mouse CD184 (CXCR4)-APC (clone L276F12, BioLegend); anti-rat CD62L-FITC (clone MEL-14, BioLegend); anti-mouse Ly6C-FITC (clone HK1.4, BioLegend). After this incubation, 500 μL of DAPI at a concentration of 1:5000 in FACS buffer was added, and the samples were centrifuged at 1800 rpm, 4°C, for 5min, with the supernatant decanted. Finally, 400 μL of counting beads (BD Trucount™, Absolute Counting Tubes, BD, 663028) resuspended in FACS buffer at a concentration of 20 beads/μL and 100μL of FACS buffer were added, resulting in a final volume of 500 μL for analysis by flow cytometry. The staining strategy for analyzing neutrophils in the ischemic hemisphere was CD45^hi^ CD11b^+^ CCR2^−^ Ly6C^−^ Ly6G^+^. To analyze neutrophils in the blood the strategy used was CD3^−^ B220^−^ Ly6C^−^ CCR2^−^ CD11b^+^. To assess neutrophils in bone marrow the strategy considered was CD11b^+^ CD115^−^ CCR2^−^ CD3^−^ B220^−^ Ly6C^−^ Ly6G^+^. Doublets and DAPI^+^ cells were excluded from analyses. Data were acquired using a FACS Canto or FACS LSRII Fortessa (BD Biosciences) and analyzed using FlowJo LLC software. Compensation was performed using single-labeled samples as reference and positive regions of staining were defined based on unstained and/or isotype-stained controls.

### Neutrophil isolation by cell sorting

Blood was extracted and lysed as described in the organ processing section for flow cytometry. Cell sorting experiments were performed using a FACS Aria cell sorter (BD Biosciences). Neutrophils were isolated by selecting TdTomato-positive cells (from Ly6G^tdTom^) in Catchup mice and as DAPI^−^ CD11B^+^ Ly6G^+^ in C57BL/6 mice.

### *Ex vivo* NET-formation assay

Neutrophils were sorted as previously described and 5×10^4^ neutrophils were plated with RPMI medium (Gibco™ RPMI 1640, HEPES) on poly-l-lysine-covered 8-well μ-Slides (Ibidi) and left 30min to adhere. Subsequently, cells were incubated for 2h with 100nM PMA or vehicle. Cells were then fixed using 4% PFA for 10min, permeabilized with PBS with 0.1% Triton X-100, 1% goat serum plus 5% BSA and stained with antibodies against cit-H3, DNA (Sytox-green, Molecular Probes) and MPO. Imaging of NETs was performed using a Zeiss LSM700 confocal microscope with 40x magnification, taking 5×5 tile-scan images with whole-slide Z-stack. We analyzed the images using Imaris (Bitplane) and identified NETs by the triple colocalization of the DNA, MPO and citH3 channels.

### Quantification of primary granules

Neutrophils sorted as previously described and plated (5×10^4^) with RPMI medium (Gibco™ RPMI 1640, HEPES) on poly-l-lysine-covered 8-well μ-Slides (Ibidi) were fixed using 4% PFA for 10 min. Afterwards, cells were stained with a biotinylated anti-MPO antibody (R&D Systems, 1:200) in PBS for 3 h at room temperature. Samples were then washed and incubated with Alexa-488 conjugated streptavidin in PBS for 1.5h at room temperature. Finally, cells were counterstained with DAPI (LifeTechnologies) to reveal nuclei. Granule images were collected using a Leica SP8 3X STED super resolution microscopy system coupled to a DMI6000 inverted microscope, with 100x magnification objective. Analysis of captured images was performed using Imaris (Bitplane) using the spots model in MPO channel.

### Neutrophil adoptive transfer experiments

Donor animals, Bmal^fl/fl^MRP8^Cre+^ and Bmal^fl/fl^MRP8^Cre-^ were injected intraperitoneally with 5 mg/kg of AMD-3100 (Sigma-Aldrich) 2 h before bleeding as described ^6,11^. A control group of mice received similar volume of saline alone at the same time points. Following intracardiac bleeding of the donors, 20μL of blood was withdrawn to assess the effectiveness of the AMD treatment and estimate the number of neutrophils injected into the recipients. The extracted blood was then distributed by 150μL intravenously administration among different naive Ly6G^tdTom^ recipient animals, 30min before starting MCAO surgery. After the surgeries, recipient animals received a second dose of 100μL of blood from their corresponding donor 5 h post-surgery.

### Neutrophil elastase activity assay

Neutrophil elastase activity in plasma was measured using a commercially available kit (Neutrophil Elastase Activity Assay kit, Fluorometric, NAK246-1KT, Sigma-Aldrich) according to the manufacturer’s instructions. We analyzed 50μl of plasma, with an excitation wavelength of 390nm and emission wavelength of 510nm in a Fluoroskan Ascent plate reader (Thermo Labsystems). Standard curve goodness-of-fit had an R2 of 0.99 in linear regression.

### Histology and tissue preparation

Mice were anesthetized and perfused intracardially with phosphate buffer (pH 7.4). The brain was removed, post-fixed with PFA overnight and cryoprotected in 30% sucrose. Finally, the brains were frozen using isopentane to proceed with sectioning on a microtome (Leica SM2000R). The sections were cut at a thickness of 30μm and stored in a cryoprotective solution, resulting in a total of 10 rostrocaudal series of sections per animal.

### Immunofluorescence in tissue

Immunofluorescence was performed on free-floating brain sections that were incubated overnight at 4°C with the following primary antibodies: anti mouse histone-3 citrulline (1:400, Abcam), anti-mouse/human myeloperoxidase (MPO) (1:200, R&D Systems), anti-mouse pan laminin biotinylated (1/250, Novus Biologicals), anti-mouse Ym1 (1/100, StemCell) and anti-mouse CD41 (1/200, Bioscience). The secondary antibodies used were anti-rabbit Alexa Fluor 647 (1:500, Thermo Fisher Scientific), anti-rat Alexa Fluor 647 (1:500, Invitrogen), streptavidin Alexa Fluor 488 (1:500, Thermo Fisher Scientific), streptavidin Alexa Fluor 405 (1:500, Thermo Fisher Scientific) and lectin 488-conjugated (1/100, Vector laboratories). Controls performed in parallel without primary antibodies showed very low levels of nonspecific staining. In some conditions, sections were incubated with DAPI (1:500, Life Technologies). Sections were then mounted on gelatin-coated slides using Aqua-Plus Mount medium (18606-100, Polysciences).

### Image acquisition, processing and quantification in tissue

Acquisitions were performed with a laser-scanning confocal imaging system (Zeiss LSM700 in CNIC). Image quantification and analysis was performed with ImageJ and Imaris.

*For NETosis analysis* a 25x objective with immersion oil was used. MPO^+^, H3C^+^, and neutrophil TdTomato^+^ were counted in confocal Z-stack images for which 3-5 serial sections (30μm) per animal spaced 300 μm apart were analyzed. Co-localization of these markers was considered positive for NETosis ^17,23^.

*For ex vivo NETosis images,* a 10x objective was used. DNA, H3C, and MPO were counted separately. Areas where the three markers co-localized or where they did not co-localize but neutrophil DNA expanded, adopting a cloud-like morphology, were considered positive for NETosis.

### Cerebral blood flow assessment

Blood perfusion was evaluated by laser speckle contrast imaging (LSCI) (RFLSI-ZW, RWD). Animals were anesthetized as described in the ischemia models section and positioned in such a way that blood flow could be recorded in two regions: the lateral area of the skull where the MCA is located, and the superior area of the skull to record cerebral blood flow through the skull, after the overlying skin had been removed. These measurements were taken before performing the craniotomy and then for 5 min, after the craniotomy. Additionally, the superior area of the calvaria was recorded for 30 min following artery occlusion, while the MCA area was recorded for 5 min. Finally, the last blood flow measurements were taken 12 h after the occlusion, with 5-minute recordings in both aforementioned areas. Blood flow was measured with the software RFLSI Analysis.02.00.31.29469.

### Neutrophil depletion

To deplete neutrophils at ZT5, 2 groups of mice were injected with either mouse anti-neutrophil antibody (anti-mouse Ly-6G 1A8 clone, which is specific to neutrophils within myeloid cells, BE0075, BioXCel; 0,4mg per day i.p.) or control isotype (rat IgG2A, BE0089, BioXCell) daily for 4 days, starting 24 h before surgery. The anti-Ly6G antibody used has been previously demonstrated to induce neutropenia with minimal changes in other blood cell populations^22^.

### *In vivo* treatments

*Chloramidine administration:* Chloramidine (Cayman Chemical Company, VITRO) was dissolved in DMSO and saline and a dosage of 50mg/kg was intravenously administered 10 min before and after MCAO. A second group of mice received similar volume of PBS alone at the same time points as control group. *DNase-I treatment:* DNase-I (recombinant human DNase-I protein) 10μg was administered intravenously, followed by an intraperitoneal injection of 50μg/250μL of the same compound 3 and 12 h after occlusion^17^.

### Assessment of neurological deficits by the footprint test

The gait pattern in mice was assessed through footprint test as described.^36^ Briefly, the fore and hind feet of the mice were painted with blue and green nontoxic paints, respectively. Animals were then allowed to freely walk along a narrow corridor (50×10×10 cm). Fresh sheet of white paper was placed on the floor of the runway for each run. The footprint patterns were analyzed for four step parameters (all measured in cm). Forelimb stride length was measured as the average distance of forward movement between each stride.

### Single cell RNA sequencing (scRNA-sequencing) of sorted tissue neutrophils

Brain neutrophils (CD11b^+^ Ly6G^+^ DAPI^−^) were isolated using a FACS Aria Fusion sorter (BD Bioscience), as detailed in the gating strategy flow-chart (Fig. 4 and Suppl. fig. 4), collected into 0.04% BSA/PBS and loaded into a BD Rhapsody cartridge. Pools of cells ten mice were created for each brain group. Up to 60.000 cells were loaded into a Rhapsody Single Cell Analysis System cartridge. Cell captures and cDNA synthesis were performed according to manufacturer’s instructions. The pool of all samples of labelled Sample Tags cells was checked for viability and cell concentration using the Countess III cell counter (Thermofisher). The average size of the libraries was calculated using the 2100 Bioanalyzer (Agilent) and the concentration was determined using the Qubit® fluorometer (Thermofisher). Finally, libraries were combined and sequenced together in a paired end run (60×42) using a NextSeq 2000 system (Illumina) and a P2 flowcell. Output files were processed with NextSeq 1000/2000 Control Software Suite v1.4.1. FastQ files for each sample were obtained using BCL Convert v3.6.3 software (Illumina). NGS experiments were performed in the Genomics Unit of the CNIC.

Sequence alignment, unique molecular identified (UMI) quantification, and single-cell identification were performed using the Rhapsody Whole Transcriptome Analysis pipeline (version 1.12.1), based on the mouse GRCm39 genome assembly and Ensembl genebuild (version 108). Single cells were filtered and clustered using scuttle (v1.10.3)^37^ and Seurat (v4.40)^38^ R packages. Cells were filtered using a sequencing depth between 1500 and 20,000, a minimum of 800 detected genes, a mitochondrial content below 10%, a gene expression complexity (percentage of reads from the top 50 genes) below 50%, a hemoglobin gene set expression below 1% and a HTO content above 50 UMIs in the hashtags. Doublets were identified using scDblFinder (v1.14.0)^39^ and were removed. Mitochondrial and hemoglobin genes were exluded for downstream analysis. At the end of the filtering process, a total of 6522 cells were retained.

Counts were normalized, and the most variable 1000 genes were identified using VST. All samples were integrated using Seurat’s IntegrateData function with RPCA method. Clustering and dimensionality reduction of these cells were performed using *15* principal components. Cluster markers and differential expression analysis between conditions have been performed using Wilcox algorithm, testing only genes detected in more than 30% of cells for markers and 10% for differential expression on any cluster. Functional analysis was performed using enrichR (v.3.2)^40^ to identify enriched pathways between conditions.

### Statistical analysis of preclinical data

For the statistical analysis of the data, Prism 9 software (GraphPad Software Inc.) was used. To compare two groups, a non-parametric Mann-Whitney test was employed. A one-way ANOVA was performed to assess differences in infarct volume among the ZT levels. Post-hoc comparisons were conducted using the Tukey HSD test to determine significant pairwise differences. For the comparison of four groups, analyzing the influence of two variables, a two-way ANOVA test with the corresponding Bonferroni post hoc test was used. Diurnal oscillations of infarct size in Fig. 1 were assessed using Cosinor analysis, which provided estimates for MESOR, amplitude, acrophase, and p-values to determine significant diurnal patterns (p<0.05). All statistical data were represented as the mean ± SEM, and in all studies, tests were considered significant when p<0.05. Excluded values from the study were identified and removed using various outlier detection criteria. The Grubbs’ test (α=0.05) was applied to exclude extreme values when only one value was considered an outlier. Additionally, the ROUT method (Q=1%) was used, and in certain cases, exclusion was based on the interquartile range (IQR), excluding values lower or higher than 1.5 times the IQR when the sample size was between 3 and 5.

**Figure 1.**
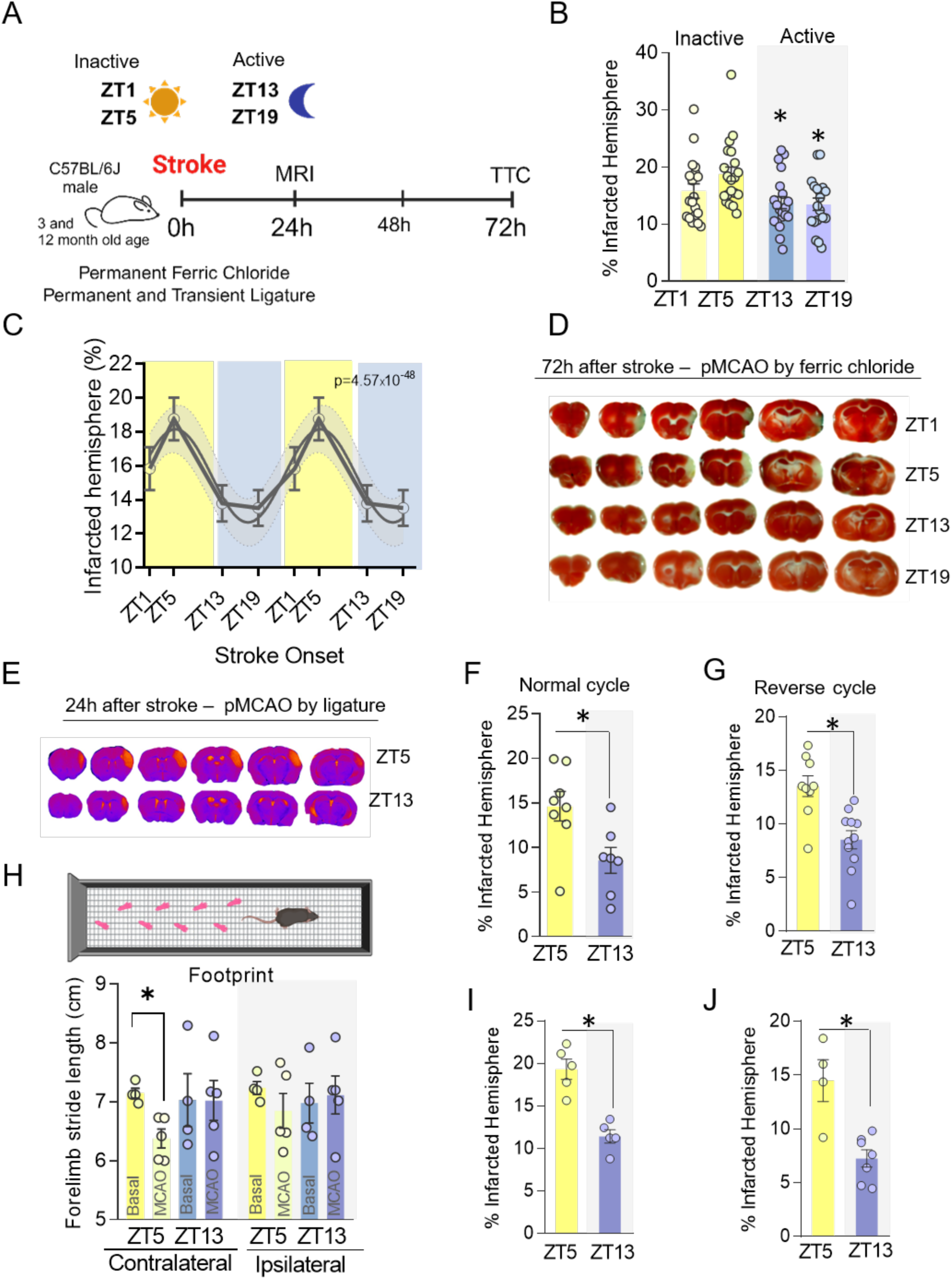
Circadian rhythms influence stroke outcome. (A) Experimental design for infarct volume determination based on activity phase (inactive., i.e. ZT5 and ZT1 or active., i.e. ZT13 and ZT19) using MRI and TTC techniques at 24h and 72h, respectively, in 3– and 12-month-old male mice in various distal ischemia models (ferric chloride and permanent or transient MCAO by ligature). (B) Percentage of infarcted hemisphere measured by TTC at 72h post-ischemia in a permanent ischemia model induced by a 20% ferric chloride application, with surgeries performed during both the inactive (ZT1 and ZT5) and the active phases (ZT13 and ZT19).*p<0.05 vs ZT5. (C) Diurnal oscillations of infarct size were assessed using COSINOR analysis. (D) Representative images of brain slices stained with TTC, measured at 72h post-ischemia in a ferric chloride model performed at ZT1, ZT5, ZT13, and ZT19, respectively. (E) Representative images of pMCAO by ligature 24 hours after stroke both at ZT5 and ZT13. Percentage of infarcted hemisphere measured by MRI 24 hours after ischemia in a permanent ligation-induced ischemia model, conducted during both the inactive phase (ZT5) and the active phase (ZT13) under a normal cycle (F) or using an inverted light cycle (G). (H) Quantification of the footprint motor test conducted between the cohort of animals operated at ZT5 and those operated at ZT13. Representation of forelimb stride length, both ipsilateral and contralateral, in centimeters, both at baseline and after ischemia for animals operated at ZT5 and ZT13. (I) Percentage of infarcted hemisphere measured by MRI 24h after ischemia in a permanent ligature-induced ischemia model, conducted during both ZT5 and ZT13 in 15-month-old animals. (J) Percentage of infarcted hemisphere measured by MRI 24 hours after ischemia in a tMCAO model by ligature, conducted at ZT5 and ZT13. Data, shown as mean ± SEM, were compared by Mann-Whitney t-test for comparing two groups (F, G-I), non-parametric Kruskal-Wallis test followed by Dunńs (B), or 2-way ANOVA followed by Bonferroni (J). *p<0.05.

### Clinical study design and patient cohort

This study included a total of 540 patients admitted to the stroke unit of Hospital 12 de Octubre (Madrid, Spain) between January 2018 and July 2024. All patients met the general inclusion and exclusion criteria detailed below.

#### Inclusion Criteria

Patients aged over 18 years with an ischemic stroke, previously independent (pre-morbid modified Rankin Scale –mRS-< 2), and admitted within six hours of symptom onset or after a wake-up stroke.

#### Exclusion Criteria

Patients with transient ischemic attacks (TIA), lacunar strokes, intracranial bleeding secondary to trauma or subarachnoid hemorrhage, or with a history of stroke, acute myocardial infarction, severe systemic infection, or major surgery within the last three months were excluded. Additional exclusion criteria included active severe systemic inflammatory disease, pregnancy, or puerperium.

#### Circadian rhythm subcohort

For the circadian rhythm analysis, a subset of 377 patients was selected from the main cohort based on the additional inclusion criterion of a witnessed stroke with a known symptom onset time. This subcohort, therefore, comprised patients with an ischemic stroke of known onset. Data on both the time of symptom onset and time of hospital admission were recorded for each patient to allow precise circadian analysis.

#### Collateral circulation subcohort

A subcohort of 310 patients from the main cohort was used to study collateral circulation. This subcohort included patients with both wake-up strokes and those with unknown symptom onset times to maximize sample size. Collateral circulation was assessed by neurologists using cerebral angiography imaging, such as computed tomography angiography (CTA) or magnetic resonance angiography (MRA). The ASITN/SIR scale (American Society of Interventional and Therapeutic Neuroradiology/Society of Interventional Radiology), which categorizes collateral circulation into five grades (0: no collaterals; 1–4: increasing degrees of collateral supply), was employed. For analysis, collateral circulation was categorized as “poor” (grades 0–2) or “good” (grades 3–4).

#### Data collection and laboratory variables

Data on age, gender, cardiovascular risk factors (CVRF), prior medication, and baseline stroke severity (National Institutes of Health Stroke Scale, NIHSS) were collected for all patients. Laboratory data, including biochemistry, cell counts (lymphocytes, leukocytes, neutrophils, platelets, mean platelet volume), and coagulation parameters, were obtained from the blood samples. Additionally, data regarding patient management, NIHSS scores at the stroke unit, infarct volumes, and 90-day mRS outcomes were retrospectively retrieved from hospital discharge and follow-up records. Neurologists measured the infarct volume for each patient using computed tomography (CT) and magnetic resonance imaging (MRI) scans obtained within the first 48 hours post-stroke. The infarct volumes were assessed in a double-blind manner to ensure objectivity and reduce assessment bias.

#### Blood sample collection and biomarker quantification

Whole blood treated with EDTA was collected upon admission and again 12–24 hours after admission while in the stroke unit. Plasma was extracted from these samples for quantification of neutrophil-specific elastase (NE), myeloperoxidase (MPO), and soluble CD40 ligand (sCD40L). Quantification was performed using commercially available enzyme-linked immunosorbent assay (ELISA) kits following the manufacturers’ instructions: Human PMN Elastase ELISA kit (Abcam, Cambridge, MA, USA), Human MPO ELISA kit (R&D Systems, Minneapolis, MN, USA), and Human CD40L (Soluble) ELISA Kit, Extra Sensitive (Thermo Fisher Scientific, Waltham, MA, USA).

#### Statistical analysis of clinical data

All analyses were conducted using the R statistical computing environment (version 4.3.0; R Foundation for Statistical Computing, Vienna, Austria) and additional specialized packages (ggplot2, dplyr, tableone, broom, ggpubr, and cosinor (version 1.2.3)). Missing values were analyzed as ‘missing’ without imputation due to the substantial proportion of missing data in certain post-stroke variables, which violated the assumptions required for robust and unbiased imputation methods. For variables with non-normal distributions, such as neutrophil elastase and myeloperoxidase (MPO), logarithmic transformations were applied to normalize the data. Normality was reassessed post-transformation using visual (QQ-plots) and statistical methods. Circadian rhythm analysis was performed using the cosinor package to model diurnal variations in biomarkers, NIHSS scores, infarct volumes, and hematological cell counts (lymphocytes, leukocytes, neutrophils, and platelets). The analysis provided estimates of MESOR (midline estimating statistic of rhythm), amplitude, and acrophase for each variable. Amplitude was of particular interest, as it reflects the extent of variation across the circadian cycle. A significant amplitude (p<0.05) indicates a meaningful fluctuation in levels over 24 hours, which may reveal underlying circadian regulation mechanisms relevant to ischemic stroke outcomes. In the collateral circulation study, comparisons were made between “poor” and “good” collateral groups. Parametric or non-parametric tests were used as appropriate for each variable after applying an interquartile range (IQR) filter to identify and exclude outliers; differential significance was set at a p-value<0.05. Sensitivity analyses confirmed that the exclusion of outliers did not alter the robustness of the results. Potential confounders were assessed using statistical tests, and no significant imbalances were detected between groups.

#### Sample size justification

This study is pioneering in evaluating diurnal patterns of immunothrombosis biomarkers, NIHSS scores, infarct volumes, and hematological cell counts in an Iberian cohort of patients with ischemic stroke. Due to the absence of prior studies in our target population using these specific markers and variables, a formal sample size calculation based on previous estimates was not feasible. However, we established a statistical significance level of α = 0.05 and a statistical power of 80% to detect clinically relevant differences. To enhance the sample size and improve statistical power in the study of collateral circulation and circadian patterns, we included patients with wake-up strokes and those with unknown symptom onset times. This decision was made because the evaluation and collection of collateral circulation grades and certain clinical parameters were not performed in all patients of the study. Although the inclusion of these patients may introduce some variability, we performed sensitivity analyses to ensure that this variability did not bias the results. This approach was deemed necessary to achieve a sufficiently large and representative sample size for robust statistical analyses.

#### Ethical approval and informed consent

This study was approved by the Institutional Review Board (IRB) of Hospital 12 de Octubre. Informed consent (CEIM 60/204) was obtained from all patients or their legal representatives prior to inclusion in the study, in compliance with institutional and ethical guidelines for human research.

## RESULTS

### CIRCADIAN RHYTHMS INFLUENCE STROKE OUTCOME

To investigate whether stroke outcome is time-of-day dependent, we first used a permanent thrombotic model caused by the topic application of ferric chloride in the middle cerebral artery (MCA). We operated mice at four different times across the day in both inactive (ZT1 and ZT5) and active (ZT13 and ZT19) phases, and quantified infarct volume 72h after stroke by using TTC (Fig. 1A-D). As observed in Fig. 1B-C, infarct size displayed marked circadian variations, with larger infarcts when occlusion was performed at daytime (at ZT5, *i.e.,* 5 h after light onset in a 12 h:12 h light:dark environment) compared to all other times in the active phase (*i.e.,* ZT13 and ZT19). To investigate the validity of these time-of-day differences (*i.e*., inactive vs. active) in stroke outcome in mice, we used a different permanent middle cerebral artery occlusion (pMCAO) model caused by the occlusion of the MCA with a ligature, at both ZT5 vs. ZT13. By using this model, in both normal and reversed 12-hour light-dark cycle (*i.e.,* the usual light phase, corresponding to the day, was shifted to the normal dark period and the other way around), we confirmed that 24h after stroke mice subjected to surgery at ZT5 displayed worse outcome as shown by higher infarct size and neurological deficits in the footprint test than those in ZT13 group (Fig. 1E-H and Supplementary Table S1). This effect of the diurnal time in stroke size was maintained in aged, 15-month-old mice (Fig. 1I). Finally, these differences in infarct size between ZT5 and ZT13 were still present in a transient MCAO model (Fig. 1J). These data demonstrate stroke-induced damage to be highly time-of-day dependent, indicating the existence of mechanisms modulated by circadian rhythms that controlled the severity of ischemic stroke.

### CIRCADIAN-DEPENDENT ISCHEMIC INJURY IS DETERMINED BY NEUTROPHILS

The number of circulating leukocytes oscillates in blood, peaking during the inactive phase in mouse^10,41,42^. Specifically, the number of circulating neutrophils peaks at around ZT5, and with low numbers by ZT13 (13h after lights on)^10^. Neutrophil size, complexity, and surface marker expression also exhibit diurnal changes, a phenomenon associated with activation and referred to as “aging”^6,10,43^. Since neutrophils are the first immune cells responding and mediating damage early after stroke onset, we tested the possibility that the circadian control of neutrophil activation was an underlying mechanism controlling the variations in stroke damage. For that, we first depleted neutrophils by treating mice with an anti-Ly6G specific antibody and evaluated the infarct size 48h after stroke in mice subjected to surgery at ZT5 and ZT13 (Fig. 2A). Neutrophil counts after 48h after stroke were lower after depletion at both times (Fig. 2B), although depletion was not complete. Importantly, antibody-mediated neutrophil depletion reduced infarct size at ZT5 compared to the isotype-treated group and led to the loss of the circadian pattern in infarct size (Fig. 2C). However, no differences were observed between isotype-treated or neutrophil-depleted ischemic mice at ZT13. These data suggest that neutrophils are responsible for at least part of the ischemic injury at ZT5 but not at ZT13. Thus, we decided to check whether time-of-day variation in stroke outcome was dependent not only on neutrophil numbers as demonstrated by the depletion model, but also on the circadian phenotype of neutrophils. For that, we performed stroke surgery at ZT5 and ZT13 and evaluated infarct size in mice in which the neutrophil clock was altered by genetic strategies (Fig. 2D-E). The neutrophil clock is governed by the key components Bmal1 and CXCR4 (Fig. 2D). BMAL1 (also known as ARNLT) can directly bind to the promoter region of CXCL2, which favors neutrophil aging in a CXCR2-dependent manner. In contrast, CXCR4 signaling blocks the CXCR2-induced aging process of neutrophils^27^. We generated neutrophil-specific BMAL– and CXCR4-deficient mice, referred to as BMAL^Neu^ and CXCR4^Neu^, by crossing Bmal1^f/f^ and CXCR4^f/f^ mice^44,45^ with mice expressing Cre recombinase in neutrophils (Mrp8-CRE)^46^. We found that, in both BMAL^Neu^ and CXCR4^Neu^ mice, ischemic damage became arrhythmic as circadian differences in infarct size between ZT5 and ZT13 were completely abolished (Fig. 2F). Interestingly, despite substantial differences in the number of brain infiltrated neutrophils among WT, BMAL^Neu^ and CXCR4^Neu^ mice, no significant differences in infarct volume were observed (Fig. 2F-G). This suggest that both neutrophil numbers and their circadian-associated phenotypes are critical factors determining ischemic damage. To further corroborate this, we performed adoptive transfer experiments with blood leukocytes isolated, as previously described^11^, from either BMAL^Neu^ mice or from mice treated with the CXCR4 antagonist AMD3100 (AMD mice), that promotes neutrophil mobilization from the bone marrow (BM) into the circulation^47^. Catchup mice, which express the red fluorescent protein tdTomato on neutrophils, were used as recipients^28^. These mice were injected intravenously, 30 minutes before and 4h after stroke at ZT5, with blood leukocytes from either WT, AMD or BMAL^Neu^ mice. 24h later, ischemic lesion and number of brain infiltrated neutrophils were analyzed (Fig. 2H-I). The transfer of leukocytes from BMAL^Neu^ mice to Catchup mice resulted in larger infarcted area compared to the transfer of WT leukocytes and with a similar trend in lesion size when recipient neutrophils derived from AMD-treated mice (Fig. 2H-I). Notably, this occurred despite lower density of infiltrated neutrophils in the brain of BMAL^Neu^ mice, suggesting that these neutrophils may exhibit a more damaging phenotype (Suppl. Fig. 1). Therefore, all these results suggest that neutrophils at the time of stroke onset are important players for determining the size of circadian ischemic injury where not only the number of neutrophils but also the phenotype might contribute to ischemic damage.

**Figure 2.**
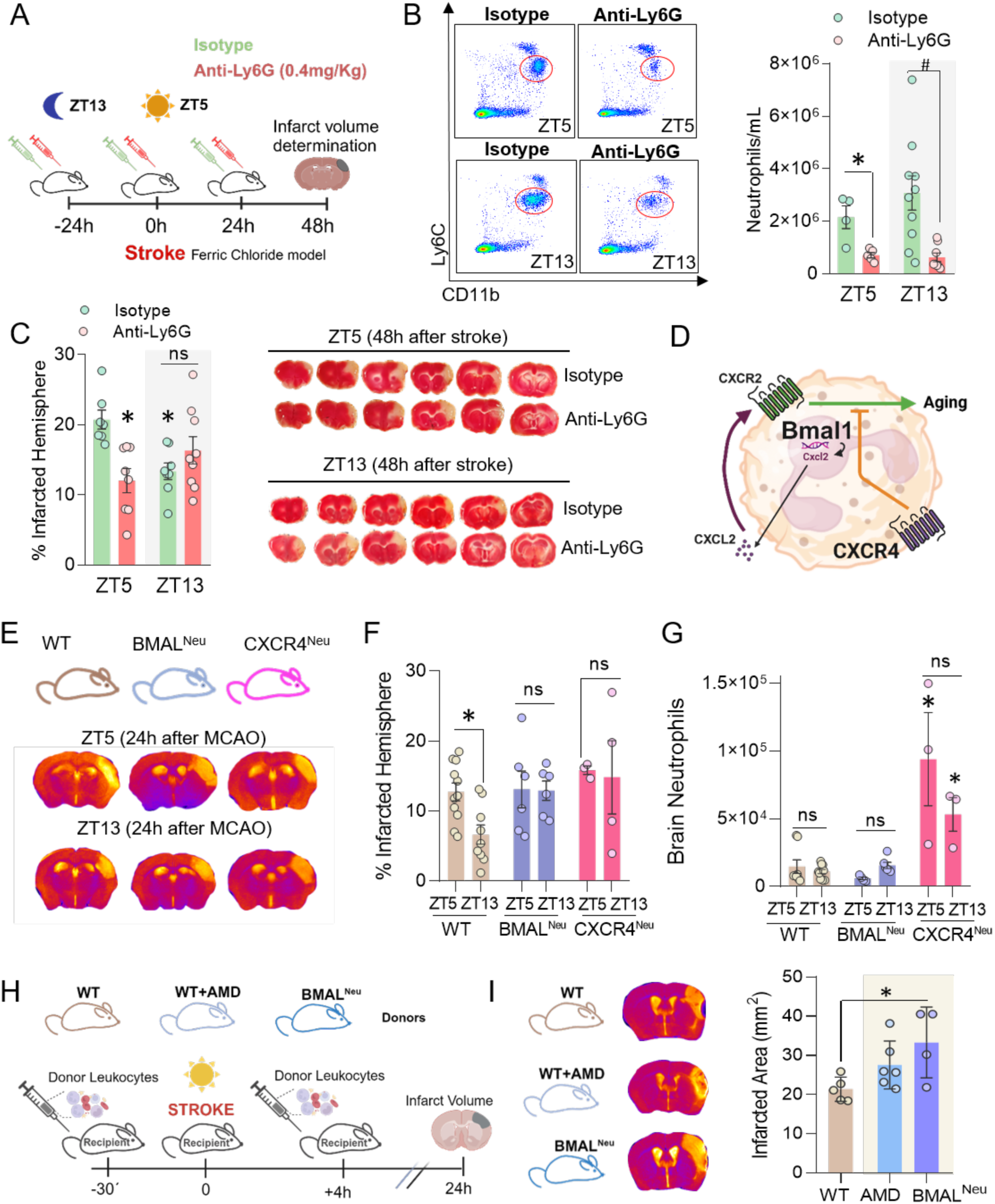
The size of ischemic injury is determined by the neutrophil circadian clock. (A) Experimental design for neutrophil inhibition through the administration of anti-Ly6G antibody (0.4 mg/kg) in a permanent ischemia model induced by ferric chloride, followed by infarct volume determination 48 hours after ischemia using TTC. (B) Left: representative images of flow cytometry plots from blood extracted 48h after ischemia at both ZT5 and ZT13, verifying the effective depletion of neutrophils in blood. The neutrophil population was gated as DAPI^−^, Ly6C^+^, and CD11b^+^ cells. Right: neutrophil quantification per ml of blood, 48h after ischemia at ZT5 and ZT13 in both isotype and anti-Ly6G-injected groups. (C) Quantification of the percentage of infarcted hemisphere after the administration of anti-Ly6G antibody in a permanent ischemia model performed at both ZT5 and ZT13, assessed 48 hours later by TTC staining (left). Representative images of TTC-stained brains (right). (D) Schematic representation of circadian aging. Bmal1 would promote the transcription of Cxcl2 and its subsequent protein translation which, once secreted from the cell, acts on the CXCR2 receptor, promoting neutrophil aging. (E) Representative MRI images of pMCAO by ligature at ZT5 and ZT13 in WT, BMAL^neu^ and CXCR4^neu^ animals 24h after ischemia. (F) Percentage of infarcted hemisphere at ZT5 and ZT13 in WT, BMAL^neu^ and CXCR4^neu^ mice. (G) Quantification of the total number of neutrophils in brain 24h after stroke in WT, BMAL^neu^ and CXCR4^Neu^ mice at both ZT5 and ZT13. (H) Experimental design for transfer of blood leukocytes from WT, WT+AMD treatment and BMAL^Neu^ donors to ischemic mice 30’ before and 4h after stroke. Infarct volume was evaluated 24h after stroke by MRI. (I) Representative images of MRI images from WT, WT with AMD treatment and BMAL^Neu^ mice 24h after stroke (left). Quantification of infarct area in WT, WT with AMD treatment and BMAL^Neu^ animals 24 hours after stroke by MRI. Data, expressed as mean ± SEM, were compared by 2-way ANOVA followed by Bonferroni (B-C and F-G) or by non-parametric Kruskal-Wallis test followed by Dunńs (I). * and # p <0.05. In (C), *p<0.05 vs ZT5.

### DAYTIME NEUTROPHILS IMPAIR CEREBRAL BLOOD FLOW RECOVERY AFTER STROKE

The circadian clock regulates neutrophil migration and infiltration throughout the body in different tissues, effectively controlling their number at specific sites and at different times of the day^27,48,49^. Since, at least partly, neutrophils mediate tissue damage after stroke by infiltrating the ischemic brain^50–54^, we next evaluated neutrophil numbers by flow cytometry in both the circulation and the brain at different times after stroke onset (5, 12 and 24h). For that, Catchup mice were subjected to stroke at ZT5 and ZT13 and neutrophils in the ischemic brain were analyzed (Fig. 3A-B). Our data show that the dynamics of neutrophils in blood (Suppl. Fig. 2) or the ischemic brain (Fig. 3B-C) were different depending on the time of the day at which mice had the ischemic event. A trend toward higher numbers of neutrophils was observed at early time points (5h after stroke) in the ischemic brain of mice subjected to surgery at ZT5 compared to ZT13 while the pattern of infiltration was inverted at latter times points (24h after stroke). In fact, 24h after stroke onset, brain neutrophil density was higher at ZT13 when compared to ZT5 further supporting the differential temporal pattern of infiltration. These distinct neutrophil migration patterns into the brain could explain the differential contribution of neutrophils at ZT5 vs ZT13, namely the impact of neutrophil depletion on infarct size at each time (Fig. 2C). Consistent with this notion, we found that the infarcted area positively correlated with the number of neutrophils in the infarcted brain at ZT5, in agreement with previous studies^51,52,55^; in contrast, no correlation was found between neutrophils and infarct size when ischemia was induced at ZT13 (Fig. 3D), highlighting the differential contribution of neutrophils to stroke at different day times. In fact, we found that, in our model, infarct volumes grew from 5h to 24h only in the ZT5 ischemic mice, while infarct growth was marginal at ZT13 (Fig. 3E).

**Figure 3.**
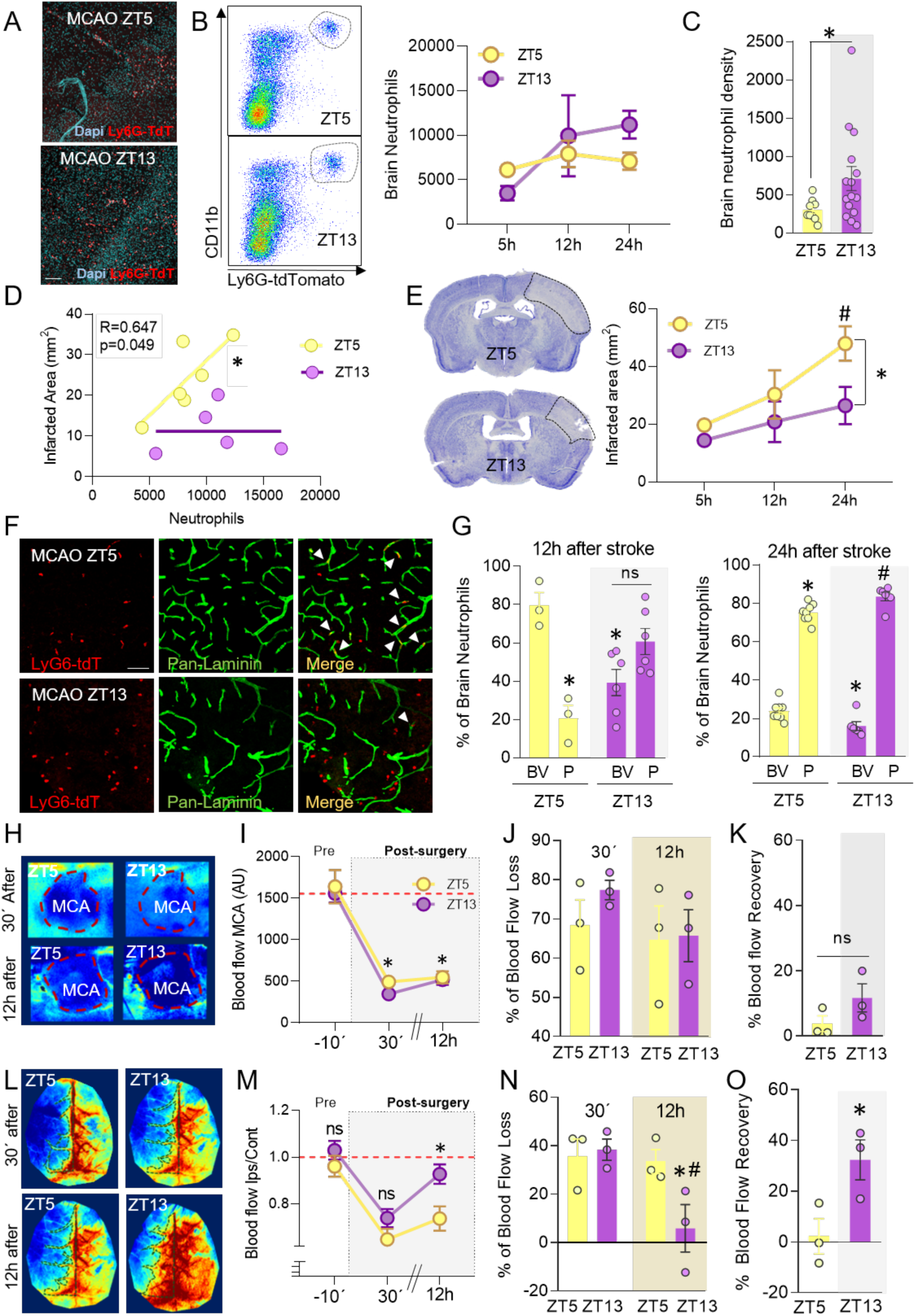
Neutrophils impair cerebral blood flow recovery after stroke in a circadian time-dependent manner. (A) Representative pictures of brain infiltrated neutrophils from Catchup mice (Ly6G-TdTomato neutrophils) in combination with DAPI (blue). Scale bar=50µm. (B) Representative flow cytometry dot plots of MCAO brains at both ZT5 and ZT13, 24h after ischemia, with neutrophils gated as DAPI^−^, CD45^+^, CD11b^+^, and Ly6G-tdTomato^+^ (left). Quantification of neutrophil count in MCAO brains at 5, 12, and 24 hours after ischemia at both ZT5 and ZT13 in a permanent ischemia model induced by ligature (right). N=3-14 mice/group. (C) Density of brain neutrophils (neutrophils/mm^2^) 24h after stroke in ZT5 and ZT13 mice. (D) Correlation between infarct area measured by MRI and neutrophil count in MCAO brains 24h after MCAO at both ZT5 and ZT13. (E) Quantification of infarct area after pMCAO induced by ligature at 5, 12, and 24 hours after ischemia using Nissl staining. Representative Nissl-stained sections 24h after stroke are shown in the left. N=3-4 mice/group. (F) Representative images of brain immunofluorescence 12h after pMCAO by ligature model, with ischemia occurring at ZT5 and ZT13, respectively. Neutrophils are marked in red by TdTomato expression, and blood vessels (BVs) are labeled in green with pan-laminin. Scale bar=50µm. (G) Percentage of total brain neutrophils located in blood vessels (BV) or parenchyma (P) 12h (left) and 24h (right) after ischemia at both ZT5 and ZT13. (H) Representative laser speckle images of the middle cerebral artery (MCA) region in a pMCAO model induced by ferric chloride, taken 30’ (top) and 12h after ischemia induction (bottom) at both ZT5 and ZT13. (I) Quantification of blood flow in the MCA region before (–10’) and 30’ and 12h after surgery, expressed as arbitrary units, at both ZT5 and ZT13. (J) Percentage of blood flow loss 30’ after ischemia compared to the pre-surgery time point at both ZT5 and ZT13 in the MCA region. (K) Percentage of blood flow recovery 12h after ischemia compared to the pre-surgery time point (–10’) at both ZT5 and ZT13 in the MCA region. (L) Representative images of both hemispheres at ZT5 and ZT13, 30’and 12h after ischemia induction. (M) Quantification of blood flow in the ipsilateral hemisphere normalized by the flow in the contralateral hemisphere before (–10’) and 30’ and 12h after surgery, expressed as arbitrary units at both ZT5 and ZT13. (N) Percentage of blood flow loss 30’ and 12h after ischemia compared to the pre-surgery time point at both ZT5 and ZT13. (O) Percentage of blood flow recovery 12h after ischemia compared to the pre-surgery time point (–10’) at both ZT5 and ZT13 in the ipsilesional hemisphere. Data, represented as mean ± SEM, were compared by Mann-Whitney t-test for comparing two groups (C, K and O), or 2-way ANOVA followed by Bonferroni (B, E, G, I-J and M-N). * and # p <0.05. In (G-), *p<0.05 vs ZT5 BV and #p<0.05 vs. ZT13 BV.

Once in the brain, neutrophils can damage tissues by releasing destructive proteolytic enzymes. In addition, they may obstruct brain capillaries, contributing to the no-reflow phenomenon and impeding full microcirculatory reperfusion^56–60^. We therefore evaluated neutrophil distribution among different compartments including their location inside blood vessels (BV) or into the brain parenchyma (P), at different times after stroke onset (i.e., 5, 12 and 24h) in mice subjected to stroke at ZT5 and ZT13. As expected, the number of neutrophils located in brain parenchyma after ischemia induced at ZT5 or ZT13 increased with time, being significantly higher at 24h compared to 5h, a time at which almost all neutrophils were located inside BVs (Suppl. Fig. 3), and to 12h after stroke (Fig. 3F-G). Evaluation of the percentage of neutrophils in both compartments, the parenchyma or inside BVs, at different time points after stroke demonstrated that neutrophil location exhibited a time-of-day dependency: 12h after stroke, neutrophils predominated in BVs in mice subjected to surgery at ZT5; in contrast, ZT13-operated mice displayed a substantial percentage of neutrophils in the parenchyma. This effect was transient since at 24h the circadian differences in distribution disappeared (Fig. 3F-G). This differential neutrophil location suggests that the detrimental actions of neutrophils at ZT5 might occur inside blood vessels. In fact, morphological analysis of brain neutrophils 5h after stroke, a time when all neutrophils are still inside BVs in both ZT5 and ZT13 groups, show that ZT5 neutrophils display an elongated morphology inside BVs compared to the rounded one observed at ZT13 (Suppl. Fig. 3). This elongation may reflect active responses to inflammatory signals in the nearby tissues^61–63^ and suggests that ZT5 neutrophils react earlier to brain damage. We postulated that the larger size and elongated morphology of ZT5 neutrophils might block brain capillaries, contributing to vascular stalling and avoiding proper cerebral perfusion and collateral circulation. To test this possibility, we evaluated cerebral blood flow (CBF) in mice 12h after surgery at ZT5 or ZT13 in the ferric chloride model by using laser speckle contrast imaging (LSCI) (Fig. 3H-O). At the MCA area, we observed a marked reduction in CBF at the time of occlusion and 12h later (Fig. 3H-K), indicating that differences in the infarct volume between ZT5 and ZT13 are not due to spontaneous recanalization of the MCA. Importantly, we also assessed CBF in the ipsilateral hemisphere 12h after MCA occlusion, as a critical time point and region for evaluating ischemic progression and potential recovery at both ZT5 and ZT13, vs. the contralateral CBF (Fig. 3L-O). Our results demonstrate a partial recovery of CBF in the ipsilesional cortical area in the ZT13 group, suggesting improvement in microvascular perfusion during this circadian phase. In contrast, the ZT5 group did not exhibit recovery in the cortical ipsilateral blood flow. Collectively, these data suggest that the timing of MCA occlusion relative to the circadian cycle might influence the degree of vascular recovery in ischemic and peri-ischemic regions, and that this process is neutrophil-dependent.

### CIRCADIAN CONTROL OF NEUTROPHIL TRANSCRIPTION AFTER STROKE

Our findings indicate that neutrophils influenced by circadian rhythms play distinct roles in ischemic stroke outcomes. To check whether circadian rhythms induce distinct neutrophil programs during stroke, we next investigated the transcriptional signature of neutrophils located in the ischemic brain (Fig. 4A). We performed scRNA-seq analysis on sorted CD11b^+^ Ly6G^+^ neutrophils from the ischemic tissue, 12h after stroke in both ZT5 and ZT13 ischemic mice. After a quality assessment (Suppl. Fig. 4), we obtained 3351 high-quality cells. These cells featured expression of neutrophil-specific genes such as S100A8, S100A9 and CSF3R and displayed low expression of the monocytic-specific CCR2 and CSFR1 (Fig. 4B). Unsupervised clustering partitioned brain neutrophils into 8 clusters (Fig. 4C-E) which display a substantial differential gene expression (Suppl. Table S2). In fact, we identified 5052 differentially expressed genes (DEGs) with specific marker genes for each of these 9 brain neutrophil clusters (BNc) like BNc0 (G0S2, Selenon, Tinf2, Slc2a3), BNc1 (*Ftl1, Fnip2, Hmox1, Gclm*), BNc2 (*Cd177, Cd47, Cstdc4, Xbp1*), BNc3 (*Pglyrp1, Rflnb, Mmp8, Retnlg*), BNc4 (*Isg15, Ifi204, Rsad2, Slfn4*) and BNc5 (*Fgl2, Fyb, Gm2a, Myadm*) (Fig. 4E and Suppl. Table S2), being the most represented clusters in the brain those from BNc0 to BNc5 (Fig. 4D). The smaller clusters BNc6 and BNc8 (*Cx3cr1, Ctss, Hexb, C1qc*) probably comprise a monocyte-related subpopulation given the high expression of *Csfr1, Ccr2 and Cx3cr1* while the cluster BNc7 (*Camp, Lyz2, Ngp, Chil3*) seems to represent the most immature subpopulation of neutrophils in the brain after stroke displaying the lowest maturation score, the highest bone marrow (BM) proximity score and the lowest neutrophil aging score (Suppl. Fig. 4 and Suppl. Table S2). We next evaluated the transcriptional profile of brain neutrophils for different core and effector neutrophil functions (Suppl. Fig. 5 and Suppl. Table S2) and found that each neutrophil subcluster displayed different scores for chemotaxis, activation, ROS production and even granule biosynthesis. We next checked whether ischemic stroke at different day times modulates the presence of the different neutrophil programs in the brain. As observed in Fig. 4F-H, the percentage of cells in each cluster dramatically fluctuates with the time of the day wherein neutrophils allocated in BNc2 and BNc4 belonged mainly to the ZT13 ischemic group (Fig. 4F-G). In particular, the BNc4 subset was highest at ZT13 and only represented 3.2% of all brain neutrophils at ZT5. In contrast, cluster BNc1 (and, in a smaller degree, the BNc3) was the most represented one in the ischemic brain when the stroke was induced at midday, with an almost 2-fold increase compared to ZT13 (Fig. 4H).

Differential gene expression and gene ontology analyses (Fig. 4J-M, Suppl. Fig. 6 and Suppl. Tables S3-4) showed that DEGs in neutrophils of BNc1 were preferentially implicated in regulating the pro-inflammatory response after stroke, by mediating immune effector processes like phagocytosis, autophagy, neutrophil activation and degranulation, and by the activation of canonical pro-inflammatory-related pathways such as TLR4 signaling, TGF-β and NF-kappa B, supporting that the BNc1 neutrophil subset is one of the subpopulations mediating damage after stroke, probably mediating the release of proteases and enzymes that mediate tissue destruction. Supporting this, DEGs in this BNc1 cluster also pointed toward a positive regulation of necrosis and apoptosis (Fig. 4I), further supporting the detrimental action of neutrophils from BNc1. BNc3 showed high expression of ribosomal genes (such as *Rps12, Rpl3 and Rpn2*) and may represent a highly plastic neutrophil phenotype that may eventually transition to other neutrophil phenotypes, as described previously in different contexts^64,65^ (Fig. 4E and Suppl. Tables S3-4). In contrast, analysis of DEGs from BNc4 indicated that this cluster responds to *interferon* (IFN) *type I* and *II* as evidenced by gene ontology analysis and the IFN score (Fig. 4J-K). Interestingly, type I IFNs have been found to trigger NET formation^66–69^. In fact, BNc4 displays one of the highest NETosis scores compared to other clusters (Suppl. Fig. 5), suggesting that neutrophils in this subset could be implicated in NETs formation, thus enhancing neutrophil-induced vascular stalling. Consistently, ZT5 neutrophils located in the cluster BNc4 displayed higher NETosis score than ZT13 ones and higher NADPH oxidase score (Fig. 4L), suggesting that neutrophils in the ZT5 ischemic brain undergo accelerated NETosis, a notion consistent with reduced numbers of neutrophils in the ZT5 ischemic brain (Fig. 3A-C). In addition, and in agreement with their differential location (Fig. 3G), neutrophils at ZT13 also displayed a significant upregulation of migration and/or extravasation related genes (like *Myo1d*, *CD47, Cd177, Lcn2)* which could explain the preferential presence of ZT13 neutrophils in the brain parenchyma (Suppl. Fig. 7 and Suppl. Tables S3-4). On the contrary, genes upregulated in ZT5 are preferentially involved in oxidative stress, iron homeostasis, different metabolic processes and even genes related to apoptosis and ferroptosis^70^ (Suppl. Fig. 7 and Suppl. Tables S3-4). These data suggest that, in experimental stroke, brain-localized neutrophils display distinct transcriptional signatures, including induction of apoptosis and NET formation, that are influenced by circadian rhythms and may impact stroke outcomes.

### CIRCADIAN MODULATION OF NEUTROPHIL FUNCTION PROMOTES NET FORMATION AFTER STROKE

Signaling via type I IFN in neutrophils has been associated with the formation of NETs^66–68^. Decreased BNc4, IFN-associated neutrophils in the brain of ischemic ZT5 mice made us hypothesize that their low levels at this time could be the result of increased neutrophil destruction by NETosis. We asked whether the differential capacity to form NETs of neutrophils at ZT5 could explain the differences in the number and subsets of infiltrated neutrophils, CBF recovery, and stroke outcome in a time-of-day dependent manner. To address this, we compared the *ex vivo* capacity of neutrophils isolated 12h after surgery at ZT5 or at ZT13. We stimulated these neutrophils with phorbol 12-myristate 13-acetate (PMA), a NET-inducing compound, and then we analyzed for the presence of NETs by colocalization of extracellular DNA (Sytox), myeloperoxidase (MPO) and citrullinated histone 3 (cit-H3) (Fig. 5).

**Figure 4.**
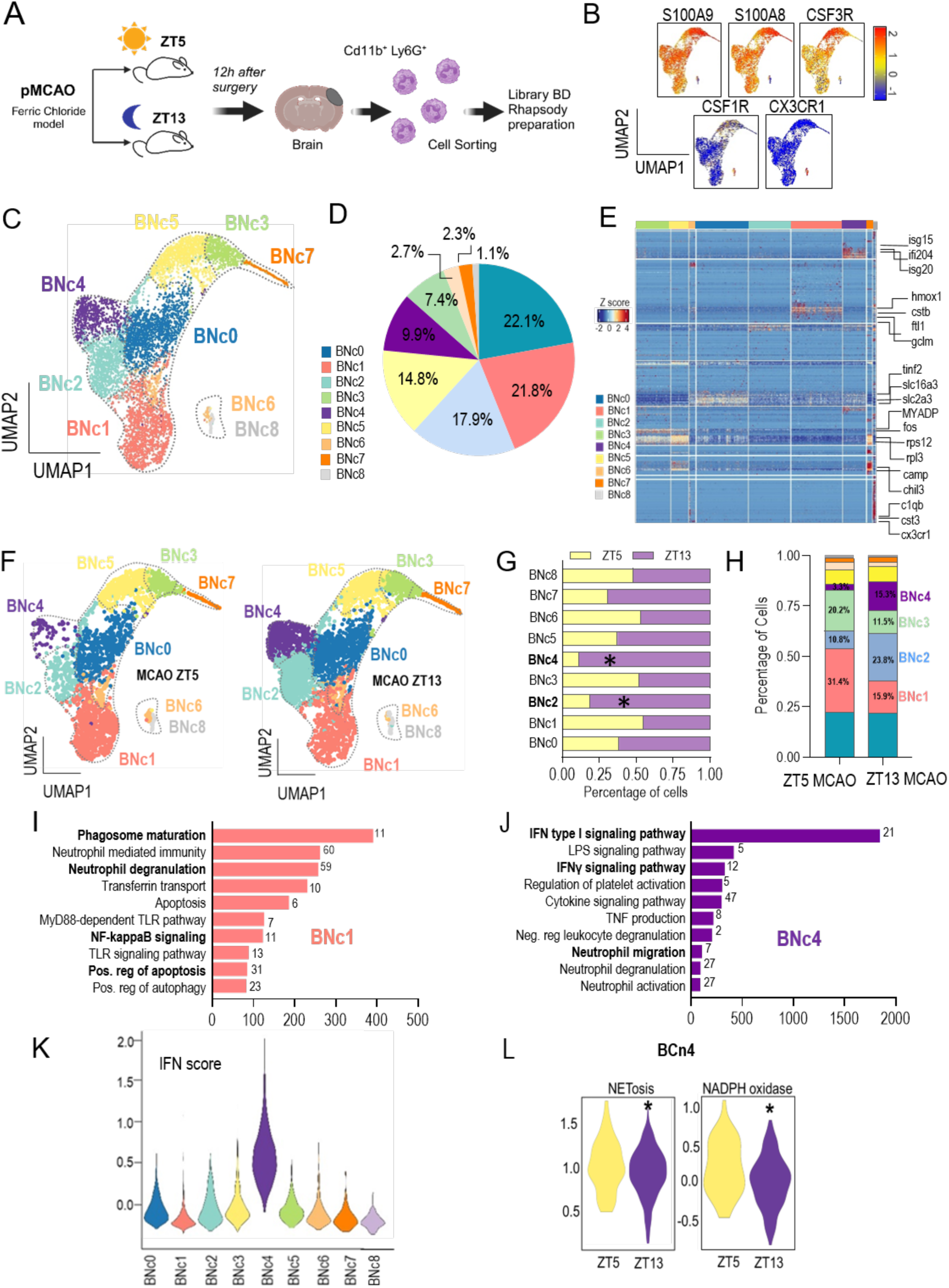
Circadian rhythms promote cerebral neutrophil heterogeneity after stroke. (A) Experimental design for scRNA-seq of brain neutrophils after stroke in mice subjected to ischemia at ZT5 or at ZT13. (B) UMAP of the 3351 brain neutrophils obtained by scRNA-seq of sorted CD45^+^CD11b^+^Ly6G^+^ cells in the ischemic cortex 12h after stroke from ZT5 and ZT13 ischemic mice, featuring the expression of neutrophil-specific genes such as S100A9, S100A8 and Csf3r and also showing low levels of expression of monocyte markers CCR2 and Csf1r. (C) UMAP of 3351 neutrophils from the brain after ischemia colored by inferred cluster identity from BNc0 to BNc8. (D) Pie chart showing the percentage of cells in each of the 9 brain neutrophil clusters. (E) Heatmap showing the row-scaled expression of the 20 highest DEGs for each of neutrophil subsets found in brain after stroke. (F) UMAPs of neutrophils colored by inferred cluster identity (from cluster BNc0 to BNc8), separated for ZT5 and ZT13 ischemic mice. (G) Graphs showing the percentage of cells from ZT5 and ZT13 in each neutrophil cluster. (H) Comparison of brain neutrophil composition between ZT5 and ZT13 ischemic mice showing the relative proportion of cells in each neutrophil cluster. (I-J) Gene ontology analysis of DEGs for the clusters BNc1 (I) and BNc4 (J). The graph shows selected gene ontology terms with Benjamini–Hochberg-corrected p-values < 0.05 and ordered by combined score. (K-L) Violin plot of interferon scores for each cluster (K) and NETosis and NADPH oxidase scores for BNc4 (L).

**Figure 5.**
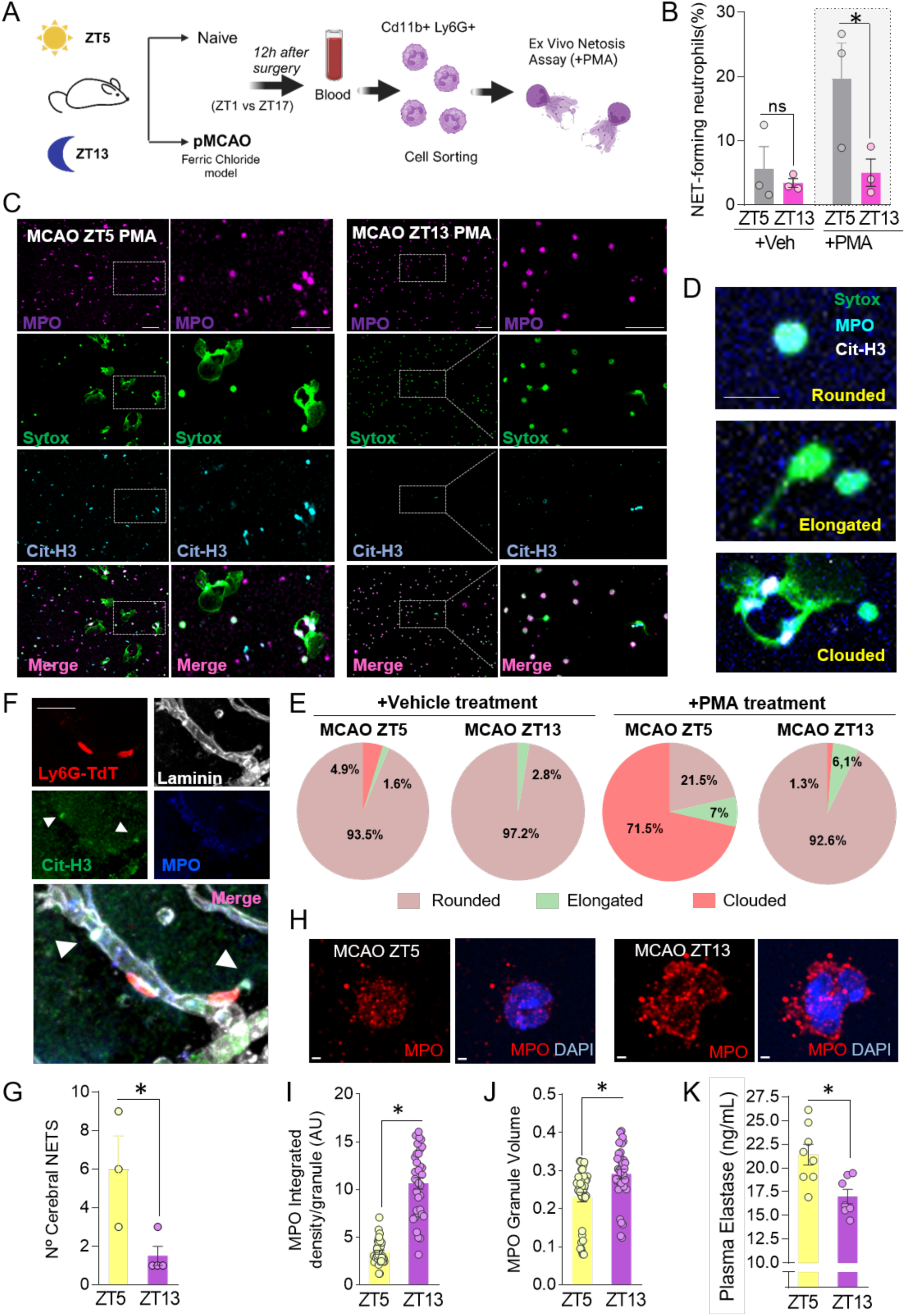
NETs promote circadian microthrombosis after stroke. (A) Experimental design for the *ex vivo* NETosis assay. Neutrophil were isolated from naive and ferric chloride-induced MCAO animals 12h after stroke and then treated with vehicle or PMA stimulation for *ex vivo* NETosis assay. (B-C) Percentage of NET-forming neutrophils in ZT5 and ZT13 MCAO animals 12h after stroke in vehicle and PMA conditions. In (C), representative images of *ex vivo* NETosis assay immunofluorescence staining in ZT5 and ZT13 MCAO animals. MPO granules are in pink, DNA is green stained by sytox green and Cit-H3 is in cyan. Scale bar=50µm. (D) Representative images of NETs morphologies (rounded, elongated and clouded) observed in *ex vivo* NETosis. Scale bar=10µm. (E) Pie charts showing the quantification of the different NETs morphologies in ZT5 and ZT13 naïve and MCAO animals upon vehicle and PMA treatment. (F) Representative images of NETosis (white arrow) in the cerebral tissue in ZT5 MCAO animals 12h after stroke. Neutrophils are in red (LyG-TdTomato), blood vessels are in white marked with pan-laminin, Cit-H3 is in green and MPO in blue. Scale bar= 50µm. (G) Numbers of cerebral NETs in ZT5 and ZT13 MCAO animals 12h after stroke. (H) Representative images of immunofluorescence staining of neutrophils isolated at 12 hours after MCAO. MPO granules are marked in red, and nuclei are stained with DAPI in blue. Scale bar=1µm. (I-J) Quantification of the MPO integrated density per granule (I) and MPO volume of neutrophil granules (J) in MCAO animals 12h post-ischemia. (K) Plasma elastase concentration (ng/ml) in ZT5 and ZT13 MCAO animals 12 hours after stroke. Data, represented as mean ± SEM, were compared by Mann-Whitney t-test for comparing two groups (C, K and O), or 2-way ANOVA followed by Bonferroni (B, E, G, I-J and M-N). *p <0.05.

We found that neutrophils isolated from the infarcted brain at ZT5 formed more NETs than those at ZT13 (Fig. 5B-C and Suppl. Fig. 8). Based on their morphological features, NETs may be classified into three subtypes according to their progression: the round ones, in which all three markers colocalize within the neutrophil^71,72^; elongated NETs, which illustrate the initiation of DNA and granular content release; and the ones with cloud morphology, as the final phase where the contents of the neutrophil assume a network shape due to the absence of elements in the surrounding environment that would limit its expansion (Fig. 5D-E). Interestingly, NETs after stroke at ZT5 preferentially displayed a clouded morphology compared to the elongated or the rounded NETs observed in neutrophils from ZT13 ischemic mice, indicating not only augmented NETs forming capacity but also an accelerated NET formation at ZT5 (Fig. 5E), and suggesting increased predisposition of these cells to release NETs. The higher potential of neutrophils from ischemic ZT5 mice to form NETs was also corroborated by the presence of NETs in the ischemic brain 12h after stroke (Fig. 5F-G). The capacity to release NETs by neutrophils has been shown to correlate with granule content^11^. In this context, neutrophil proteases, such as MPO, which are stored in azurophilic granules, play a crucial role in degrading nuclear histones and facilitating chromatin decondensation during the process of NET formation. Consistently, confocal microscopy of Ly6G^+^ blood neutrophils 12h after stroke (Fig. 5H-J and Suppl. Fig. 9) revealed that neutrophils from ischemic ZT13 mice display higher MPO intensity per granule and higher granule size than ZT5 ones indicating that MPO content and NET-forming capacity are inversely correlated in blood neutrophils from ischemic mice. In fact, the MPO^+^ granule content in neutrophils was also inversely correlated with plasma elastase levels (another protein stored in azurophilic granules) (Fig. 5K), suggesting the release of granule content into blood and an increased degranulation potential of neutrophils from ZT5 ischemic mice. Taken together, our findings indicate that NETosis is the circadian-dependent mechanism by which neutrophils drive vascular stalling following stroke.

### INHIBITION OF NET FORMATION ABOLISHES CIRCADIAN DIFFERENCES IN STROKE OUTCOME

To investigate the influence of NETosis on infarct volume after ischemic stroke, we next inhibited NET formation by two distinct strategies: a pharmacological approach and a genetic intervention, both of which target the peptidyl arginine deiminase 4 (PAD4), a key enzyme for histone citrullination during NET formation (Fig. 6). We first administered chloramidine, a pharmacological inhibitor of PAD4, and measured infarct volumes 24h post-surgery (Fig. 6A). NET inhibition by chloramidine produced a significant reduction in the lesion size compared to the vehicle-treated group at ZT5, but not at ZT13 (Fig. 6B). Similarly, PAD4^−/−^ mice showed a significant reduction in the infarct volume at ZT5 compared to control mice but not at ZT13 (Fig. 6C), supporting the time-restricted contribution of NETs in the brain damage during the inactive phase (ZT5).

**Figure 6.**
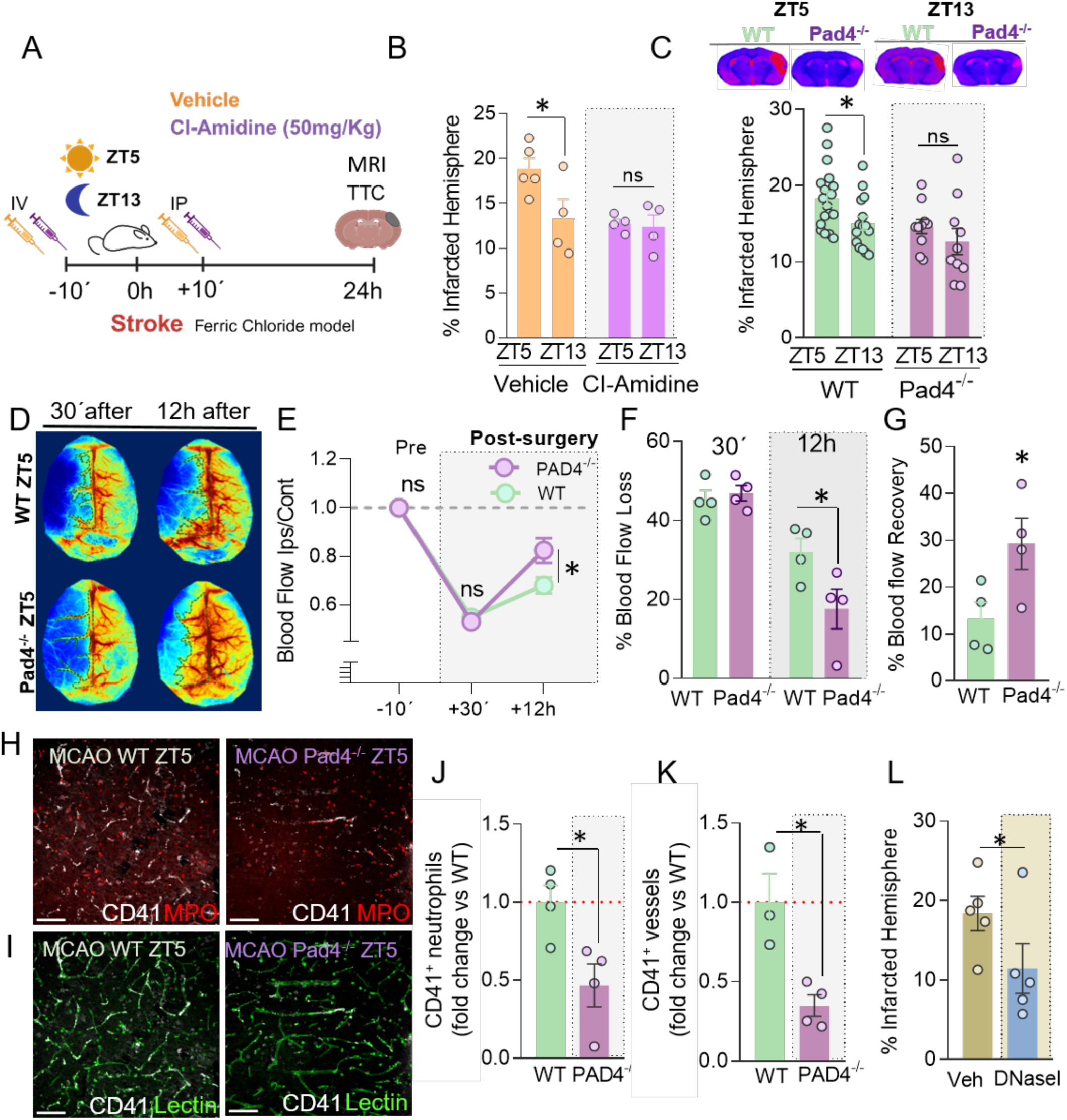
Inhibition of NETs abolished circadian-dependent differences in stroke outcome. (A) Experimental design for NETosis inhibition by chloramidine (Cl-Amidine; 50mg/kg IV and IP) in ZT5 and ZT13 pMCAO by ferric chloride model and determination of infarct size by TTC 24h after stroke. (B) Percentage of infarcted hemisphere in ZT5 and ZT13 pMCAO mice and determination of infarct size by TTC 24h after stroke. (C) Percentage of infarcted hemisphere in ZT5 and ZT13 pMCAO ferric chloride model, in WT and PAD4^−/−^ animals, and determination of infarct size by MRI 24h after stroke. Representative MRI images of brains 24h after stroke are shown in the top panel. (D) Representative images of CBF by LSI of both hemispheres at ZT5 from WT and PAD^−/−^ mice before (30’ before) and 12h after ischemia induction (12h after). (E) CBF in the ipsilateral hemisphere normalized by the flow in the contralateral hemisphere before surgery (–10’) and 30’ and 12h after surgery, in WT and PAD^−/−^ ischemic mice subjected to surgery at ZT5. (F) Percentage of CBF loss 30’ after ischemia compared to the pre-surgery time point in WT and PAD^−/−^ ischemic mice subjected to surgery at ZT5. (G) Percentage of CBF recovery 12h after ischemia (12 h) compared to the pre-surgery time point (–10’ before) in the ipsilesional cortex of WT and PAD^−/−^ ischemic mice subjected to surgery at ZT5. (H) Representative images of neutrophils and platelets in pMCAO WT and PAD4^−/−^ mouse brains 24h after stroke conducted at ZT5. MPO^+^ granules in neutrophils are in red and platelets are in white stained with CD41. (I) Representative images of cerebral tissue in ZT5 pMCAO WT and PAD4^−/−^ animals 24 hours after stroke, where blood vessels are stained for lectin (green) and platelets marked with CD41 antibody (white). Scale bar=50µm. (J) Platelet-neutrophil interactions (CD41^+^ neutrophils) at ZT5 in pMCAO PAD4^−/−^ and WT animals, expressed as fold-change vs WT. (K) BVs occluded by platelets 24h after stroke in the brain of WT and PAD^−/−^ ischemic mice subjected to surgery at ZT5, expressed as fold-change vs WT. (L) Percentage of the infarcted hemisphere in mice subjected to pMCAO at ZT5 and treated with either vehicle or DNase-I. Infarct was analyzed by MRI 24h after stroke. Data, represented as mean ± SEM, were compared by Mann-Whitney t-test for two groups (G, I and K) and 2-way ANOVA followed by Bonferroni (B-C, E-F and L). *p <0.05.

The role of NETs during ischemic stroke was also confirmed in another experimental stroke model induced by *in situ* thrombin injection (Suppl. Fig. 10). In addition, NET inhibition improved the CBF recovery after MCAO, consistent with reduced vascular stalling (Fig. 6D-K and Suppl. Fig. 11). Quantification of blood flow in this region in control and PAD4^−/−^ mice subjected to surgery at ZT5 demonstrated higher recovery of blood flow in the ipsilesional cortical area in PAD4^−/−^ mice (Fig. 6D-G). In agreement with a recovered CBF, vascular occluded segments in this cortical region by both neutrophils and platelets was also reduced in PAD4^−/−^ mice (Fig. 6H-K). Finally, the intravenous administration of DNase-I after MCAO, as a strategy for reducing NET-dependent stalling at ZT5, caused a marked reduction in infarct volume at this time (Fig. 6L).

Altogether, these data support that neutrophil stalling of the vasculature, interactions with platelets and NET release, suggestive of intravascular immunothrombosis, mediate impaired vascular perfusion and underlie circadian-dependent brain injury after stroke, posing the idea that strategies directed to modulate diurnal neutrophil reprograming and NET formation or to promote NET degradation are promising therapies for stroke patients.

### DIURNAL OSCILLATIONS IN HUMAN STROKE SEVERITY, IMMUNOTHROMBOSIS AND NET MARKERS LINKED TO COLLATERAL CIRCULATION

Finally, to explore the time-of-day dependent influence of NETosis on ischemic stroke severity in humans, we next analyzed data from 377 patients with known time of stroke onset (Fig. 7). Stroke patients had a median age of 76 years, 51.2% were female, and they displayed a median NIHSS score on admission of 10 (Suppl. Tables S5-6). Using an adjusted COSINOR model fitted to the time of admission, we found that different hematological parameters, including neutrophil and platelet counts, mean platelet volume (MPV) and plasma markers associated with immunothrombosis (sCD40L) and NET release (elastase and MPO) all displayed diurnal fluctuations, peaking at 9 and 6-7 PM, respectively (Fig. 7A, Suppl. Fig. 12 and Suppl. Table S7). Supporting the notion that NETs formation and vascular stalling influence infarct progression and stroke severity, our stroke patient cohort also displayed time-of-day oscillations in several stroke outcome parameters, including admission and 24h NIHSS score and infarct volumes, that aligned with the corresponding circadian rhythmicity observed for NET release and immunothrombosis markers (Fig. 7B and Suppl. Table S7).

**Figure 7.**
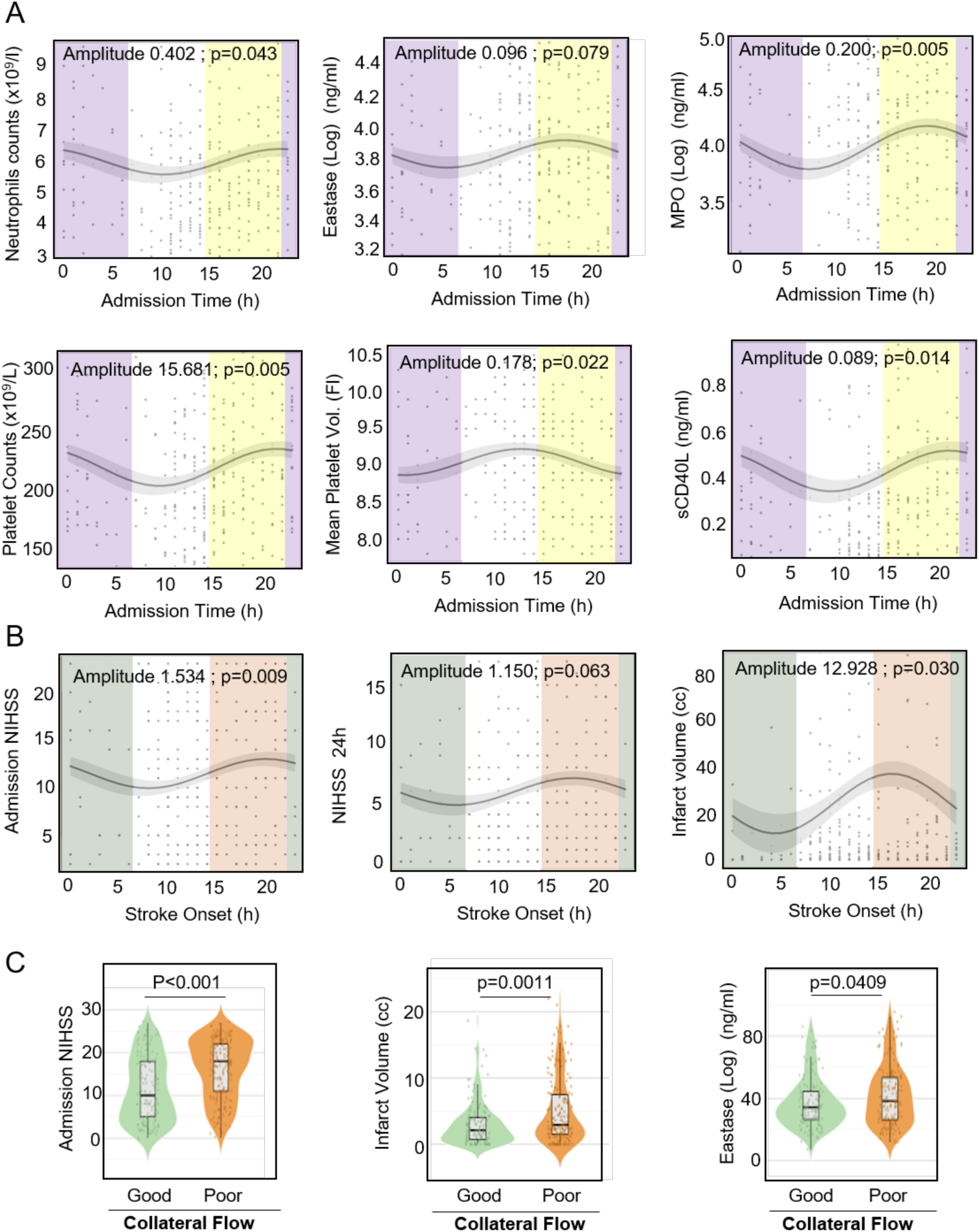
Stroke patients display diurnal oscillations in stroke severity and NET markers linked to collateral circulation. (A) Diurnal oscillations of admission hematological parameters including neutrophils and platelets counts, mean platelet volume and plasma markers related to NETosis and immunothrombosis (sCD40L, elastase, and MPO) aligned with admission time. Time of day segmentation was performed in 8-h intervals following published recommendations^98^. (B) Diurnal variations of clinical stroke severity parameters, including admission NIHSS, NIHSS at 24h and infarct volume, according to stroke onset time. Each plot presents the COSINOR-fitted curve with a shaded ribbon representing the standard error of the fit. The amplitude and p-value of each oscillation are indicated in the plot titles; rhythms with p-values below 0.05 are considered to have significant diurnal patterns. (C) Comparative analysis of clinical markers by collateral circulation status using violin plots. Violin plots illustrate the distribution of admission NIHSS, infarct volume and admission elastase in patients with good versus poor collateral circulation (n= 349). Each violin plot includes data points (jittered) and displays the median as a white diamond. Outliers were removed using the interquartile range (IQR) method, with outliers defined as values lying beyond 1.5 times the IQR from the first and third quartiles. For each marker, appropriate statistical tests were conducted based on data distribution. t-test applied to admission elastase and Wilcoxon tests applied to admission NIHSS and infarct size. *p <0.05.

Finally, given the potential relationship between collateral circulation and stroke severity, we explored whether this association also extended to markers of NETosis in a sub-cohort of 349 stroke patients. For that, violin plots were generated to illustrate the distribution of key clinical markers, including first prognostic markers (admission NIHSS and infarct volume) and markers of NETs formation (admission elastase and MPO) in patients with either good or poor collateral circulation (Fig. 7C). As expected, stroke patients with good collateral circulation showed lower infarct volume and admission NIHSS score than those with a poor collateral circulation (Fig. 7C) and, importantly and supporting our previous results, plasma levels of admission elastase were also increased in patients with a poor collateral flow strongly supporting that NETosis is also a key factor in the differential susceptibility of humans to stroke, depending on circadian rhythms.

## DISCUSSION

Ischemic stroke is a devastating vascular disease with significant mortality and morbidity. Despite decades of research into neuroprotective treatments, none has proven effective enough for clinical use^73^. Time-of-day has emerged as a significant factor underlying the translational gap in the clinical management of ischemic stroke^2,4^ so understanding how these rhythms affect stroke responses and stroke prognosis could provide valuable insights to improve therapeutic strategies for stroke patients. Here, we demonstrate, both in mice and human, that stroke severity displays diurnal circadian oscillations and identify neutrophil-mediated NETosis and vascular stalling as key contributor mechanisms accounting for the time-of-day-dependent differential severity after stroke. Therefore, strategies aimed at modulating neutrophil heterogeneity, inhibiting NETosis formation, or degrading NETs are promising targets for therapeutic intervention in stroke management. Importantly, it is vital to emphasize that, for any of these treatments to be effective, the influence of circadian rhythms must be taken into account. Identifying the optimal time of day in humans to assess the efficacy of each treatment could be crucial for maximizing their impact on cerebral perfusion in stroke but also in different pathological scenarios where circadian rhythms and inflammation play a fundamental role in the development and progression of the pathology^74–76^.

In the present work, we have used different permanent and transient mice models to evaluate the influence of circadian rhythms on stroke outcome. Our findings demonstrate that infarct size is significantly larger when the stroke occurred at ZT5, during the inactive mice phase, compared to the one in active mice period, with a clear pattern of circadian oscillations as observed by the COSINOR fitted model. This pattern was consistent across various models, including aged animals, at both normal and reverse cycles and was associated with neuromotor deficits, such as a reduction in contralateral forelimb stride length at ZT5 in the footprint test. Similar data was previously observed in both mice and rats^4,6,77,78^. In addition, our data also demonstrate that diurnal circadian oscillations in stroke severity also occur in human ischemic stroke, in agreement with our data in animal models and with previous studies showing that patients with stroke symptom onset at night, i.e., during the human inactive period, showed larger core volumes and a faster core size progression compared to patients during the human active period^79,80^. Importantly, our findings in both humans and mouse models show consistent results across various markers, aligning with the respective activity phases of each species. Although mice are nocturnal and humans are diurnal, the observed patterns in infarct volume and associated markers remain congruent when adjusted for each species active phase. This cross-species consistency underscores the relevance of circadian influences on stroke outcomes and highlights the robustness of our results across different biological rhythms.

Circadian rhythms may play a significant role in shaping the progression and outcomes of ischemic stroke by affecting critical components of the ischemic cascade. Among those mechanisms, the inflammatory response and the immune system are attractive candidates^5^. In this sense, different immune cell types operate on a circadian schedule, demonstrating 24-hour rhythmicity both under resting conditions and during periods of activation. From all these cell types, neutrophils, which are the first responders to inflammation following a stroke, display circadian variations in their numbers, phenotypes, and activation markers^10,41–43^. These fluctuations made us think that neutrophils may influence stroke outcomes depending on the time of the day. Indeed, neutrophil reduction after stroke with a specific anti-Ly6G antibody abolished circadian-dependent differences in stroke outcome. Of note, not only neutrophil numbers seem to be important: the inhibition of the neutrophil clock by using genetic strategies with specific-neutrophil deletion of either BMAL1 or CXCR4 produced a loss of circadian variation in infarct size compared to wild-type animals, supporting the important role of fluctuations in both neutrophil number and phenotype in stroke severity. This effect was confirmed by adoptive transfer experiments wherein similar numbers of BMAL1 KO transferred neutrophils promote higher infarcts than those neutrophils from WT mice, also supporting the key role of the neutrophil phenotype.

With the evidence that both neutrophil counts and phenotype shape infarct size in ischemic stroke, we sought to uncover their exact contributions. We found that the temporal pattern of neutrophil infiltration into the brain and, importantly, their location inside blood vessels (BVs) or into the brain parenchyma was quite different for stroke occurring at ZT5 vs. ZT13. In this sense, our data suggest that detrimental actions of neutrophils at ZT5 (when infarct size is higher) occur inside BVs instead of into the brain parenchyma. In fact, a positive correlation between infarct size and neutrophil counts was observed only at ZT5, suggesting a more damaging phenotype at this time. The viability of this region, in addition to being influenced by collateral flow, is vulnerable to secondary microcirculatory failure^55,81,82^. This failure is characterized by dynamic flow stalls in the capillaries, which persist in the salvageable penumbra even after reperfusion, and in which neutrophils appear to play a key role^19,83^. In this context, immunostaining revealed a greater number of intravascular neutrophils and neutrophil-platelet aggregates^61^, likely obstructing vessels at ZT5, suggesting circadian-related cerebral blood flow stalling in a neutrophil-dependent way.

Although neutrophils have been traditionally considered a detrimental cell in the context of ischemic stroke^50–53,84^, experiments involving neutrophil depletion or inhibition of their brain infiltration yield inconsistent results, suggesting a more complex role for these immune cells. Indeed, recent research has revealed that neutrophils exhibit a remarkable degree of plasticity and heterogeneity, with distinct subpopulations depending on the context^6,85–88^ that contribute differentially to both physiology and pathology. Consistently, previous studies in cerebral ischemia have demonstrated that neutrophils are not a homogeneous population in both blood and brain^21–24^. In this sense, our study provides additional evidence supporting that, upon stroke and depending on the time of the day, brain neutrophils display different phenotypes and functions that could either worsen or improve tissue damage after stroke. Of note, our data show that, upon stroke at ZT13, there is a prominent representation of neutrophil clusters which are almost absent at ZT5. In this line, the BNc4 represents an interferon (IFN)-induced neutrophil cluster that is activated or influenced by IFN signaling, particularly by both type I (like IFN-α and IFN-β) and type II IFNs (IFN-γ). These cytokines play a crucial role in the immune response, especially in the context of viral infections, inflammatory diseases and autoimmune conditions. Specifically, the response to IFN caught our attention due to its implications in NETosis^89–92^, a process whereby neutrophils release web-like structures of DNA, histones, and proteases known as neutrophil extracellular traps (NETs), that play a crucial role in immunothrombosis, inflammation and blood-brain barrier disruption^89,90,93^ and have been implicated in the pathogenesis of ischemic stroke in part by mediating stalling of brain capillaries^19,83^.

Indeed, IFN promotes NETosis by inducing the expression of specific proteins such as peptidylarginine deiminase 4 (PAD4), which is essential for citrullination of histones and subsequent NET formation; additionally, type I IFNs can modulate the activation state of neutrophils, further facilitating NETosis in response to various stimuli, including viral infections^66–69^. Surprisingly, the BNc4 neutrophil cluster is upregulated at ZT13, when infarct size is smaller. This unexpected result made us hypothesize that, given the deleterious role of NETosis previously demonstrated in the context of ischemic stroke^13,17,18,93–95^, neutrophils at ZT5 exhibit an accelerated pattern of NETosis as well as an enhanced response to IFN when compared with those neutrophils at ZT13. Congruently, we speculated that these neutrophils may not be detectable in our single-cell RNA sequencing because, 12h post-stroke, they may have undergone destruction through NETosis. Interestingly, similar results were observed in a study in which, using scRNA-seq, a decrease in neutrophil counts in the lungs of COVID-19-associated pulmonary aspergillosis patients was found that was attributed to increased neutrophil destruction *via* NETosis^96^. To assess such a possibility in our setting, we isolated blood neutrophils and evaluated their NETosis capacity. Importantly, our data show increased NETosis at ZT5 compared to ZT13 in both circulating and brain-resident neutrophils. ELISA confirmed higher plasma elastase levels, a NETosis proxy, at ZT5, and brain immunostaining revealed a greater NET burden.

Differences in stroke outcome at ZT5 vs. ZT13 could be due to reduced blood flow in ZT5 mice, which might result from NETosis-induced vascular obstruction at ZT5. Consistently, pharmacological or genetic inhibition of NET led to a reduction in infarct volume only at ZT5, again underscoring a circadian-dependent effect. Blood flow analyses in PAD4^−/−^ animals revealed improved collateral circulation at ZT5 compared to WT animals, reinforcing the role of NETs in blood flow disruption and suggesting that NETosis exacerbates microvascular stalling in a time-specific manner. Importantly, similar fluctuations in NETosis markers were observed in our human cohort that significantly align with good or poor collateral flow, then reinforcing the notion that NETs formation as an immunothrombotic mechanism hindering cerebral perfusion recovery after stroke.

Finally, the treatment with DNase-I, that degrades NETs, proved to be effective in reducing infarct volume at ZT5, supporting its potential to counteract the time-dependent damage exacerbation caused by NETosis. These findings are confirmed by recent evidence showing that DNAse-I reduces infarct volumes after in situ thromboembolic stroke, a tPA-sensitive stroke mouse mode, in the inactive but not in the active rodent phase^97^, highlighting the therapeutic relevance of DNase-I in mitigating circadian influences on ischemic damage and support targeting NETs as a strategy to improve stroke outcomes.

Collectively, our findings suggest that circadian influences on neutrophil count and phenotype and neutrophil-related functions such as NETosis play a critical role in the severity of ischemic injury. This evidence not only helps to explain the translational failures of neuroprotective therapies but also suggest that strategies targeting neutrophil function and NET formation and degradation may be more effective at specific times of the day, thus maximizing their protective effects. Our data also support the potential efficacy of personalized chronotherapy in stroke management based in specific, NETosis-inspired, plasma biomarkers. Future therapies should therefore integrate chronobiological principles to optimize patient outcomes. It is worth noting that ongoing clinical trials are investigating the efficacy and safety of several therapies, including DNase-I and its impact on various inflammatory and neurovascular pathologies. Our studies will help establish the effects of this and other strategies in a clinical context and refine the timing and conditions of its administration according to circadian rhythms. Accounting for circadian influences on neutrophils and their associated processes, such as NETosis, could offer a novel and effective strategy in developing therapeutic interventions for stroke.

## ACKNOWLEDGMENTS

We thank all members of our laboratory for insightful feedback. E. Prieto, R. Nieto, M. Vitón, N.A Muñoz and B. Álvarez-Flores for help with sorting and cytometric analyses; E. Garrido and S. Rodriguez for animal husbandry; E. Arza, O. Giménez, V. Labrador, V. Caiolfa from the Microscopy Unit of the CNIC for help with microscopy.

## DATA AND MATERIALS AVAILABILITY

The data are presented in the main manuscript and in the supplementary materials. RNA-seq data are deposited in the Genome Expression Omnibus (GEO) with accession number GSE290318.

## REFERENCES

1. O’Collins VE, Macleod MR, Donnan GA, Horky LL, van der Worp BH, Howells DW. 1,026 experimental treatments in acute stroke. Ann Neurol 2006;59:467–77.

2. Lo EH, Albers GW, Dichgans M, Donnan G, Esposito E, Foster R, Howells DW, Huang YG, Ji X, Klerman EB and others. Circadian Biology and Stroke. Stroke 2021;52:2180–2190.

3. Mergenthaler P, Balami JS, Neuhaus AA, Mottahedin A, Albers GW, Rothwell PM, Saver JL, Young ME, Buchan AM. Stroke in the Time of Circadian Medicine. Circ Res 2024;134:770–790.

4. Esposito E, Li W, T Mandeville E, Park JH, Şencan I, Guo S, Shi J, Lan J, Lee J, Hayakawa K and others. Potential circadian effects on translational failure for neuroprotection. Nature 2020;582:395–398.

5. Esposito E, Zhang F, Park JH, Mandeville ET, Li W, Cuartero MI, Lizasoaín I, Moro MA, Lo EH. Diurnal Differences in Immune Response in Brain, Blood and Spleen After Focal Cerebral Ischemia in Mice. Stroke 2022;53:e507–e511.

6. Adrover JM, Del Fresno C, Crainiciuc G, Cuartero MI, Casanova-Acebes M, Weiss LA, Huerga-Encabo H, Silvestre-Roig C, Rossaint J, Cossío I and others. A Neutrophil Timer Coordinates Immune Defense and Vascular Protection. Immunity 2019;50:390–402.e10.

7. Lembach A, Stahr A, Ali AAH, Ingenwerth M, von Gall C. Sex-Dependent Effects of Bmal1-Deficiency on Mouse Cerebral Cortex Infarction in Response to Photothrombotic Stroke. Int J Mol Sci 2018;19(10):3124.

8. Liu JA, Walker WH, Meléndez-Fernández OH, Bumgarner JR, Zhang N, Walton JC, Meares GP, DeVries AC, Nelson RJ. Dim light at night shifts microglia to a pro-inflammatory state after cerebral ischemia, altering stroke outcome in mice. Exp Neurol 2024;377:114796.

9. Kamat PK, Khan MB, Siddiqui S, Williams D, da Silva Lopes Salles E, Naeini SE, Arbab AS, Rudic DR, Baban B, Dhandapani KM and others. Time Dimension Influences Severity of Stroke and Heightened Immune Response in Mice. Transl Stroke Res 2023. Dec 13. doi: 10.1007/s12975-023-01226-5.

10. Casanova-Acebes M, Pitaval C, Weiss LA, Nombela-Arrieta C, Chèvre R, A-González N, Kunisaki Y, Zhang D, van Rooijen N, Silberstein LE and others. Rhythmic modulation of the hematopoietic niche through neutrophil clearance. Cell 2013;153:1025–35.

11. Adrover JM, Aroca-Crevillén A, Crainiciuc G, Ostos F, Rojas-Vega Y, Rubio-Ponce A, Cilloniz C, Bonzón-Kulichenko E, Calvo E, Rico D and others. Programmed ‘disarming’ of the neutrophil proteome reduces the magnitude of inflammation. Nat Immunol 2020;21:135–144.

12. Grover SP, Mackman N. Neutrophils, NETs, and immunothrombosis. Blood 2018;132:1360–1361.

13. Zhu S, Yu Y, Qu M, Qiu Z, Zhang H, Miao C, Guo K. Neutrophil extracellular traps contribute to immunothrombosis formation via the STING pathway in sepsis-associated lung injury. Cell Death Discov 2023;9(1):315.

14. Brinkmann V, Reichard U, Goosmann C, Fauler B, Uhlemann Y, Weiss DS, Weinrauch Y, Zychlinsky A. Neutrophil extracellular traps kill bacteria. Science 2004;303:1532–5.

15. Yang H, Biermann MH, Brauner JM, Liu Y, Zhao Y, Herrmann M. New Insights into Neutrophil Extracellular Traps: Mechanisms of Formation and Role in Inflammation. Front Immunol 2016;7:302.

16. Silvestre-Roig C, Braster Q, Ortega-Gomez A, Soehnlein O. Neutrophils as regulators of cardiovascular inflammation. Nat Rev Cardiol 2020;17:327–340.

17. Peña-Martínez C, Durán-Laforet V, García-Culebras A, Ostos F, Hernández-Jiménez M, Bravo-Ferrer I, Pérez-Ruiz A, Ballenilla F, Díaz-Guzmán J, Pradillo JM and others. Pharmacological Modulation of Neutrophil Extracellular Traps Reverses Thrombotic Stroke tPA (Tissue-Type Plasminogen Activator) Resistance. Stroke 2019;50:3228–3237.

18. Denorme F, Portier I, Rustad JL, Cody MJ, de Araujo CV, Hoki C, Alexander MD, Grandhi R, Dyer MR, Neal MD and others. Neutrophil extracellular traps regulate ischemic stroke brain injury. J Clin Invest 2022;132(10):e154225.

19. El Amki M, Glück C, Binder N, Middleham W, Wyss MT, Weiss T, Meister H, Luft A, Weller M, Weber B and others. Neutrophils Obstructing Brain Capillaries Are a Major Cause of No-Reflow in Ischemic Stroke. Cell Rep 2020;33:108260.

20. Aroca-Crevillén A, Vicanolo T, Ovadia S, Hidalgo A. Neutrophils in Physiology and Pathology. Annu Rev Pathol 2024;19:227–259.

21. Cuartero MI, Ballesteros I, Moraga A, Nombela F, Vivancos J, Hamilton JA, Corbi AL, Lizasoain I, Moro MA. N2 Neutrophils, Novel Players in Brain Inflammation After Stroke: Modulation by the PPARgamma Agonist Rosiglitazone. Stroke. 2013;44:3498–508.

22. García-Culebras A, Durán-Laforet V, Peña-Martínez C, Moraga A, Ballesteros I, Cuartero MI, de la Parra J, Palma-Tortosa S, Hidalgo A, Corbí AL and others. Role of TLR4 (Toll-Like Receptor 4) in N1/N2 Neutrophil Programming After Stroke. Stroke 2019;50:2922–2932.

23. Gullotta GS, De Feo D, Friebel E, Semerano A, Scotti GM, Bergamaschi A, Butti E, Brambilla E, Genchi A, Capotondo A and others. Age-induced alterations of granulopoiesis generate atypical neutrophils that aggravate stroke pathology. Nat Immunol 2023;24:925–940.

24. Garcia-Bonilla L, Shahanoor Z, Sciortino R, Nazarzoda O, Racchumi G, Iadecola C, Anrather J. Analysis of brain and blood single-cell transcriptomics in acute and subacute phases after experimental stroke. Nat Immunol 2024;25:357–370.

25. Percie du Sert N, Hurst V, Ahluwalia A, Alam S, Avey MT, Baker M, Browne WJ, Clark A, Cuthill IC, Dirnagl U and others. The ARRIVE guidelines 2.0: updated guidelines for reporting animal research. BMJ Open Sci 2020;4(1):e100115.

26. Casanova-Acebes M, Nicolás-Ávila JA, Li JL, García-Silva S, Balachander A, Rubio-Ponce A, Weiss LA, Adrover JM, Burrows K, A-González N and others. Neutrophils instruct homeostatic and pathological states in naive tissues. J Exp Med 2018;215:2778–2795.

27. Adrover JM, Del Fresno C, Crainiciuc G, Cuartero MI, Casanova-Acebes M, Weiss LA, Huerga-Encabo H, Silvestre-Roig C, Rossaint J, Cossío I and others. A Neutrophil Timer Coordinates Immune Defense and Vascular Protection. Immunity 2019;50:390–402.e10.

28. Hasenberg A, Hasenberg M, Männ L, Neumann F, Borkenstein L, Stecher M, Kraus A, Engel DR, Klingberg A, Seddigh P and others. Catchup: a mouse model for imaging-based tracking and modulation of neutrophil granulocytes. Nat Methods 2015;12:445–52.

29. Jiménez-Alcázar M, Rangaswamy C, Panda R, Bitterling J, Simsek YJ, Long AT, Bilyy R, Krenn V, Renné C, Renné T and others. Host DNases prevent vascular occlusion by neutrophil extracellular traps. Science 2017;358:1202–1206.

30. Ballesteros I, Cuartero MI, Moraga A, de la Parra J, Lizasoain I, Moro M. Stereological and flow cytometry characterization of leukocyte subpopulations in models of transient or permanent cerebral ischemia. J Vis Exp 2014 (94):52031.

31. Cuartero MI, de la Parra J, Pérez-Ruiz A, Bravo-Ferrer I, Durán-Laforet V, García-Culebras A, García-Segura JM, Dhaliwal J, Frankland PW, Lizasoain I and others. Abolition of aberrant neurogenesis ameliorates cognitive impairment after stroke in mice. J Clin Invest 2019;129:1536–1550.

32. Pradillo JM, Denes A, Greenhalgh AD, Boutin H, Drake C, McColl BW, Barton E, Proctor SD, Russell JC, Rothwell NJ and others. Delayed administration of interleukin-1 receptor antagonist reduces ischemic brain damage and inflammation in comorbid rats. J Cereb Blood Flow Metab 2012;32:1810–9.

33. Karatas H, Erdener SE, Gursoy-Ozdemir Y, Gurer G, Soylemezoglu F, Dunn AK, Dalkara T. Thrombotic distal middle cerebral artery occlusion produced by topical FeCl(3) application: a novel model suitable for intravital microscopy and thrombolysis studies. J Cereb Blood Flow Metab 2011;31:1452–60.

34. García-Yébenes I, Sobrado M, Zarruk JG, Castellanos M, Pérez de la Ossa N, Dávalos A, Serena J, Lizasoain I, Moro MA. A mouse model of hemorrhagic transformation by delayed tissue plasminogen activator administration after in situ thromboembolic stroke. Stroke 2011;42:196–203.

35. García-Culebras A, Palma-Tortosa S, Moraga A, García-Yébenes I, Durán-Laforet V, Cuartero MI, de la Parra J, Barrios-Muñoz AL, Díaz-Guzmán J, Pradillo JM and others. Toll-Like Receptor 4 Mediates Hemorrhagic Transformation After Delayed Tissue Plasminogen Activator Administration in In Situ Thromboembolic Stroke. Stroke 2017;48:1695–1699.

36. Hernández-Jiménez M, Peña-Martínez C, Godino MDC, Díaz-Guzmán J, Moro M, Lizasoain I. Test repositioning for functional assessment of neurological outcome after experimental stroke in mice. PLoS One 2017;12:e0176770.

37. McCarthy DJ, Campbell KR, Lun AT, Wills QF. Scater: pre-processing, quality control, normalization and visualization of single-cell RNA-seq data in R. Bioinformatics 2017;33:1179–1186.

38. Hao Y, Hao S, Andersen-Nissen E, Mauck WM, Zheng S, Butler A, Lee MJ, Wilk AJ, Darby C, Zager M and others. Integrated analysis of multimodal single-cell data. Cell 2021;184:3573–3587.e29.

39. Germain PL, Lun A, Garcia Meixide C, Macnair W, Robinson MD. Doublet identification in single-cell sequencing data using. F1000Res 2021;10:979.

40. Kuleshov MV, Jones MR, Rouillard AD, Fernandez NF, Duan Q, Wang Z, Koplev S, Jenkins SL, Jagodnik KM, Lachmann A and others. Enrichr: a comprehensive gene set enrichment analysis web server 2016 update. Nucleic Acids Res 2016;44(W1):W90–7.

41. Méndez-Ferrer S, Lucas D, Battista M, Frenette PS. Haematopoietic stem cell release is regulated by circadian oscillations. Nature 2008;452:442–7.

42. Scheiermann C, Kunisaki Y, Frenette PS. Circadian control of the immune system. Nat Rev Immunol 2013;13:190–8.

43. Zhang D, Chen G, Manwani D, Mortha A, Xu C, Faith JJ, Burk RD, Kunisaki Y, Jang JE, Scheiermann C and others. Neutrophil ageing is regulated by the microbiome. Nature 2015;525(7570):528-32.

44. Nie Y, Waite J, Brewer F, Sunshine MJ, Littman DR, Zou YR. The role of CXCR4 in maintaining peripheral B cell compartments and humoral immunity. J Exp Med 2004;200:1145–56.

45. Janich P, Pascual G, Merlos-Suárez A, Batlle E, Ripperger J, Albrecht U, Cheng HY, Obrietan K, Di Croce L, Benitah SA. The circadian molecular clock creates epidermal stem cell heterogeneity. Nature 2011;480:209–14.

46. Passegué E, Wagner EF, Weissman IL. JunB deficiency leads to a myeloproliferative disorder arising from hematopoietic stem cells. Cell 2004;119:431–43.

47. Devi S, Wang Y, Chew WK, Lima R, A-González N, Mattar CN, Chong SZ, Schlitzer A, Bakocevic N, Chew S and others. Neutrophil mobilization via plerixafor-mediated CXCR4 inhibition arises from lung demargination and blockade of neutrophil homing to the bone marrow. J Exp Med 2013;210:2321–36.

48. Schloss MJ, Horckmans M, Nitz K, Duchene J, Drechsler M, Bidzhekov K, Scheiermann C, Weber C, Soehnlein O, Steffens S. The time-of-day of myocardial infarction onset affects healing through oscillations in cardiac neutrophil recruitment. EMBO Mol Med 2016;8:937–48.

49. He W, Holtkamp S, Hergenhan SM, Kraus K, de Juan A, Weber J, Bradfield P, Grenier JMP, Pelletier J, Druzd D and others. Circadian Expression of Migratory Factors Establishes Lineage-Specific Signatures that Guide the Homing of Leukocyte Subsets to Tissues. Immunity 2018;49:1175–1190.e7.

50. Perez-de-Puig I, Miró-Mur F, Ferrer-Ferrer M, Gelpi E, Pedragosa J, Justicia C, Urra X, Chamorro A, Planas AM. Neutrophil recruitment to the brain in mouse and human ischemic stroke. Acta Neuropathol 2015;129:239–57.

51. Neumann J, Riek-Burchardt M, Herz J, Doeppner TR, König R, Hütten H, Etemire E, Männ L, Klingberg A, Fischer T and others. Very-late-antigen-4 (VLA-4)-mediated brain invasion by neutrophils leads to interactions with microglia, increased ischemic injury and impaired behavior in experimental stroke. Acta Neuropathol 2015;129:259–77.

52. Neumann J, Henneberg S, von Kenne S, Nolte N, Müller AJ, Schraven B, Görtler MW, Reymann KG, Gunzer M, Riek-Burchardt M. Beware the intruder: Real time observation of infiltrated neutrophils and neutrophil-Microglia interaction during stroke in vivo. PLoS One 2018;13:e0193970.

53. Kalimo H, del Zoppo GJ, Paetau A, Lindsberg PJ. Polymorphonuclear neutrophil infiltration into ischemic infarctions: myth or truth? Acta Neuropathol 2013;125:313–6.

54. Price CJ, Menon DK, Peters AM, Ballinger JR, Barber RW, Balan KK, Lynch A, Xuereb JH, Fryer T, Guadagno JV and others. Cerebral neutrophil recruitment, histology, and outcome in acute ischemic stroke: an imaging-based study. Stroke 2004;35:1659–64.

55. Jickling GC, Liu D, Ander BP, Stamova B, Zhan X, Sharp FR. Targeting neutrophils in ischemic stroke: translational insights from experimental studies. J Cereb Blood Flow Metab 2015;35:888–901.

56. Kloner RA, King KS, Harrington MG. No-reflow phenomenon in the heart and brain. Am J Physiol Heart Circ Physiol 2018;315:H550–H562.

57. Connolly ES, Winfree CJ, Springer TA, Naka Y, Liao H, Yan SD, Stern DM, Solomon RA, Gutierrez-Ramos JC, Pinsky DJ. Cerebral protection in homozygous null ICAM-1 mice after middle cerebral artery occlusion. Role of neutrophil adhesion in the pathogenesis of stroke. J Clin Invest 1996;97:209–16.

58. Ember JA, del Zoppo GJ, Mori E, Thomas WS, Copeland BR, Hugli TE. Polymorphonuclear leukocyte behavior in a nonhuman primate focal ischemia model. J Cereb Blood Flow Metab 1994;14:1046–54.

59. del Zoppo GJ, Schmid-Schönbein GW, Mori E, Copeland BR, Chang CM. Polymorphonuclear leukocytes occlude capillaries following middle cerebral artery occlusion and reperfusion in baboons. Stroke 1991;22:1276–83.

60. Mori E, del Zoppo GJ, Chambers JD, Copeland BR, Arfors KE. Inhibition of polymorphonuclear leukocyte adherence suppresses no-reflow after focal cerebral ischemia in baboons. Stroke 1992;23:712–8.

61. Sreeramkumar V, Adrover JM, Ballesteros I, Cuartero MI, Rossaint J, Bilbao I, Nácher M, Pitaval C, Radovanovic I, Fukui Y and others. Neutrophils scan for activated platelets to initiate inflammation. Science 2014;346:1234–8.

62. Phillipson M, Heit B, Colarusso P, Liu L, Ballantyne CM, Kubes P. Intraluminal crawling of neutrophils to emigration sites: a molecularly distinct process from adhesion in the recruitment cascade. J Exp Med 2006;203:2569–75.

63. Ley K, Laudanna C, Cybulsky MI, Nourshargh S. Getting to the site of inflammation: the leukocyte adhesion cascade updated. Nat Rev Immunol 2007;7:678–89.

64. Salcher S, Sturm G, Horvath L, Untergasser G, Kuempers C, Fotakis G, Panizzolo E, Martowicz A, Trebo M, Pall G and others. High-resolution single-cell atlas reveals diversity and plasticity of tissue-resident neutrophils in non-small cell lung cancer. Cancer Cell 2022;40:1503–1520.e8.

65. Goveia J, Rohlenova K, Taverna F, Treps L, Conradi LC, Pircher A, Geldhof V, de Rooij LPMH, Kalucka J, Sokol L and others. An Integrated Gene Expression Landscape Profiling Approach to Identify Lung Tumor Endothelial Cell Heterogeneity and Angiogenic Candidates. Cancer Cell 2020;37(1):21–36.e13.

66. Moreira-Teixeira L, Stimpson PJ, Stavropoulos E, Hadebe S, Chakravarty P, Ioannou M, Aramburu IV, Herbert E, Priestnall SL, Suarez-Bonnet A and others. Type I IFN exacerbates disease in tuberculosis-susceptible mice by inducing neutrophil-mediated lung inflammation and NETosis. Nat Commun 2020;11(1):5566.

67. Rocha BC, Marques PE, Leoratti FMS, Junqueira C, Pereira DB, Antonelli LRDV, Menezes GB, Golenbock DT, Gazzinelli RT. Type I Interferon Transcriptional Signature in Neutrophils and Low-Density Granulocytes Are Associated with Tissue Damage in Malaria. Cell Rep 2015;13:2829–2841.

68. Glennon-Alty L, Moots RJ, Edwards SW, Wright HL. Type I interferon regulates cytokine-delayed neutrophil apoptosis, reactive oxygen species production and chemokine expression. Clin Exp Immunol 2021;203:151–159.

69. González-Navajas JM, Lee J, David M, Raz E. Immunomodulatory functions of type I interferons. Nat Rev Immunol 2012;12:125–35.

70. Dixon SJ, Lemberg KM, Lamprecht MR, Skouta R, Zaitsev EM, Gleason CE, Patel DN, Bauer AJ, Cantley AM, Yang WS and others. Ferroptosis: an iron-dependent form of nonapoptotic cell death. Cell 2012;149:1060–72.

71. Cichon I, Santocki M, Ortmann W, Kolaczkowska E. Imaging of Neutrophils and Neutrophil Extracellular Traps (NETs) with Intravital (In Vivo) Microscopy. Methods Mol Biol 2020;2087:443–466.

72. Santocki M, Kolaczkowska E. On Neutrophil Extracellular Trap (NET) Removal: What We Know Thus Far and Why So Little. Cells 2020;9(9):2079.

73. Lyden P, Buchan A, Boltze J, Fisher M, Consortium* SX. Top Priorities for Cerebroprotective Studies-A Paradigm Shift: Report From STAIR XI. Stroke 2021;52:3063–3071.

74. Aziz IS, McMahon AM, Friedman D, Rabinovich-Nikitin I, Kirshenbaum LA, Martino TA. Circadian influence on inflammatory response during cardiovascular disease. Curr Opin Pharmacol 2021;57:60–70.

75. Thosar SS, Butler MP, Shea SA. Role of the circadian system in cardiovascular disease. J Clin Invest 2018;128:2157–2167.

76. Rana S, Prabhu SD, Young ME. Chronobiological Influence Over Cardiovascular Function: The Good, the Bad, and the Ugly. Circ Res 2020;126:258–279.

77. Mandeville ET, Li W, Quinto-Alemany D, Zhang F, Esposito E, Nakano T, Mandeville JB, Lee J, Park JH, Arai K and others. Fingolimod Does Not Reduce Infarction After Focal Cerebral Ischemia in Mice During Active or Inactive Circadian Phases. Stroke 2022;53:3741–3750.

78. Kamat PK, Khan MB, Wood K, Siddiqui S, Rudic DR, Dhandapani K, Waller J, Hess DC. Preclinical evaluation of circadian rhythm in ischemic stroke outcomes. Cond Med 2021;4:280-284.

79. Reidler P, Brehm A, Sporns PB, Burbano VG, Stueckelschweiger L, Broocks G, Liebig T, Psychogios MN, Ricke J, Dimitriadis K and others. Circadian rhythm of ischaemic core progression in human stroke. J Neurol Neurosurg Psychiatry 2023;94:70–73.

80. Ryu WS, Hong KS, Jeong SW, Park JE, Kim BJ, Kim JT, Lee KB, Park TH, Park SS, Park JM and others. Association of ischemic stroke onset time with presenting severity, acute progression, and long-term outcome: A cohort study. PLoS Med 2022;19:e1003910.

81. Iadecola C, Alexander M. Cerebral ischemia and inflammation. Curr Opin Neurol 2001;14:89–94.

82. Semerano A, Laredo C, Zhao Y, Rudilosso S, Renú A, Llull L, Amaro S, Obach V, Planas AM, Urra X and others. Leukocytes, Collateral Circulation, and Reperfusion in Ischemic Stroke Patients Treated With Mechanical Thrombectomy. Stroke 2019;50:3456–3464.

83. Erdener Ş, Tang J, Kılıç K, Postnov D, Giblin JT, Kura S, Chen IA, Vayisoğlu T, Sakadžić S, Schaffer CB and others. Dynamic capillary stalls in reperfused ischemic penumbra contribute to injury: A hyperacute role for neutrophils in persistent traffic jams. J Cereb Blood Flow Metab 2021;41:236–252.

84. Price CJ, Menon DK, Peters AM, Ballinger JR, Barber RW, Balan KK, Lynch A, Xuereb JH, Fryer T, Guadagno JV and others. Cerebral neutrophil recruitment, histology, and outcome in acute ischemic stroke: an imaging-based study. Stroke; a journal of cerebral circulation 2004;35:1659–64.

85. Ballesteros I, Rubio-Ponce A, Genua M, Lusito E, Kwok I, Fernández-Calvo G, Khoyratty TE, van Grinsven E, González-Hernández S, Nicolás-Ávila J and others. Co-option of Neutrophil Fates by Tissue Environments. Cell 2020;183:1282–1297.e18.

86. Ng LG, Ostuni R, Hidalgo A. Heterogeneity of neutrophils. Nat Rev Immunol 2019;19:255–265.

87. Xie X, Shi Q, Wu P, Zhang X, Kambara H, Su J, Yu H, Park SY, Guo R, Ren Q and others. Single-cell transcriptome profiling reveals neutrophil heterogeneity in homeostasis and infection. Nat Immunol 2020;21:1119–1133.

88. Grieshaber-Bouyer R, Radtke FA, Cunin P, Stifano G, Levescot A, Vijaykumar B, Nelson-Maney N, Blaustein RB, Monach PA, Nigrovic PA and others. The neutrotime transcriptional signature defines a single continuum of neutrophils across biological compartments. Nat Commun 2021;12:2856.

89. Bressan A, Faggin E, Donato M, Tonon L, Buso R, Nardin C, Tiepolo M, Cinetto F, Scarpa R, Agostini C and others. NETosis in Acute Thrombotic Disorders. Semin Thromb Hemost 2023;49:709–715.

90. Sharma S, Hofbauer TM, Ondracek AS, Chausheva S, Alimohammadi A, Artner T, Panzenboeck A, Rinderer J, Shafran I, Mangold A and others. Neutrophil extracellular traps promote fibrous vascular occlusions in chronic thrombosis. Blood 2021;137:1104–1116.

91. Thakur M, Junho CVC, Bernhard SM, Schindewolf M, Noels H, Döring Y. NETs-Induced Thrombosis Impacts on Cardiovascular and Chronic Kidney Disease. Circ Res 2023;132:933–949.

92. Thålin C, Hisada Y, Lundström S, Mackman N, Wallén H. Neutrophil Extracellular Traps: Villains and Targets in Arterial, Venous, and Cancer-Associated Thrombosis. Arterioscler Thromb Vasc Biol 2019;39:1724–1738.

93. Kang L, Yu H, Yang X, Zhu Y, Bai X, Wang R, Cao Y, Xu H, Luo H, Lu L and others. Neutrophil extracellular traps released by neutrophils impair revascularization and vascular remodeling after stroke. Nat Commun 2020;11:2488.

94. Novotny J, Oberdieck P, Titova A, Pelisek J, Chandraratne S, Nicol P, Hapfelmeier A, Joner M, Maegdefessel L, Poppert H and others. Thrombus NET content is associated with clinical outcome in stroke and myocardial infarction. Neurology 2020;94:e2346–e2360.

95. Datsi A, Piotrowski L, Markou M, Köster T, Kohtz I, Lang K, Plöttner S, Käfferlein HU, Pleger B, Martinez R and others. Stroke-derived neutrophils demonstrate higher formation potential and impaired resolution of CD66b + driven neutrophil extracellular traps. BMC Neurol 2022;22:186.

96. Feys S, Vanmassenhove S, Kraisin S, Yu K, Jacobs C, Boeckx B, Cambier S, Cunha C, Debaveye Y, Gonçalves SM and others. Lower respiratory tract single-cell RNA sequencing and neutrophil extracellular trap profiling of COVID-19-associated pulmonary aspergillosis: a single centre, retrospective, observational study. Lancet Microbe 2024;5:e247–e260.

97. Di G, Vázquez-Reyes S, Díaz B, Peña-Martinez C, García-Culebras A, Cuartero MI, Moraga A, Pradillo JM, Esposito E, Lo EH and others. Daytime DNase-I Administration Protects Mice From Ischemic Stroke Without Inducing Bleeding or tPA-Induced Hemorrhagic Transformation, Even With Aspirin Pretreatment. Stroke 2025;56:527–532.

98. Saver JL, Klerman EB, Buchan AM, Calleja P, Lizasoain I, Bahr-Hosseini M, Lee S, Liebeskind DS, Mergenthaler P, Mun KT, and others; Leducq Consortium International pour la Recherche Circadienne sur l’AVC Effects in Stroke (CIRCA). Consensus Recommendations for Standardized Data Elements, Scales, and Time Segmentations in Studies of Human Circadian/Diurnal Biology and Stroke. Stroke. 2023;54:1943–1949.

