## Supplemental Material for "Circadian Modulation of Neutrophil Function Determines Collateral Perfusion and Outcome After Ischemic Stroke"

Supplementary Figures 1 to 12

Supplementary Tables S1 to S7

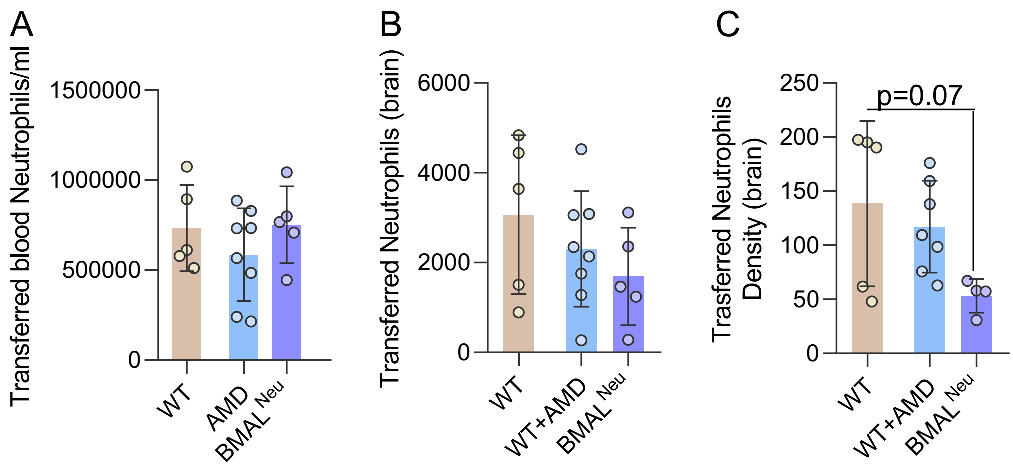


D

| Donor | 30 min before stroke | | 4 h after stroke | |
|---|---|---|---|---|
| | **Blood volume (µl)** | **Neutrophils (Mean)** | **Blood volume (µl)** | **Neutrophils (Mean)** |
| WT | 120 | 68,992 | 100 | 82,814 |
| WT + AMD | 120 | 969,422 | 100 | 571,275 |
| BMAL$^{neu}$ | 150 | 757,000 | 100 | 627,822 |

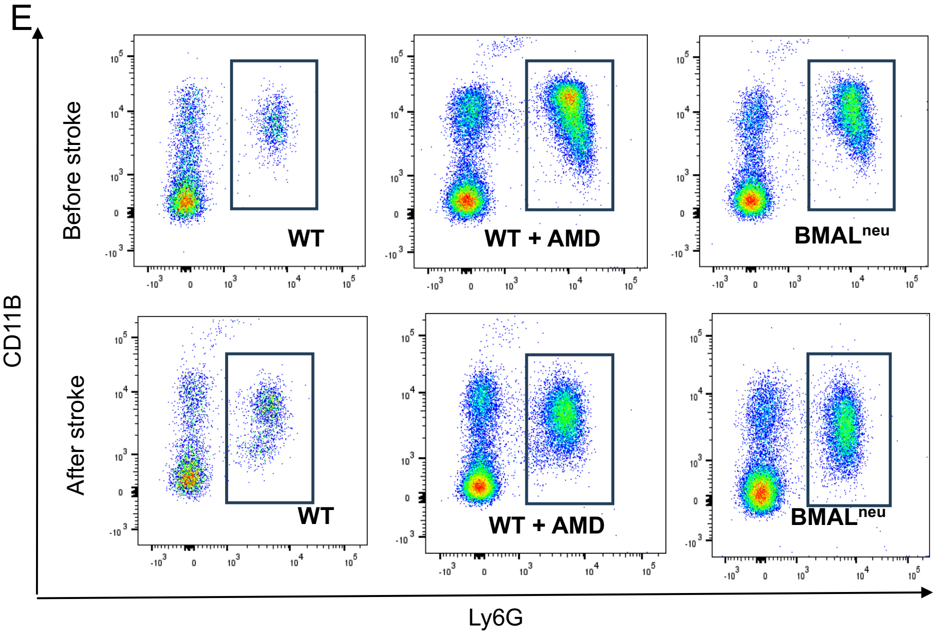


**Supplementary Figure 1. Quantification and flow cytometry analysis of transferred neutrophils.** (A) Quantification of transferred blood neutrophils/ml, (B) transferred brain neutrophils and (C) transferred neutrophils density from donors in receptor animals 24 hours after ischemia in WT group, AMD+WT group and BMAL$^{neu}$ group. A two-way ANOVA test was performed followed by Bonferroni post-hoc test, considering a p-value < 0.05 as significant. (D) Summary table with the number of neutrophils transferred from each group in the two injections performed for each mouse (before stroke and after stroke). (E) Representative cytometry dot plots of blood neutrophils from each of the donor groups (WT, WT + AMD and BMAL$^{neu}$ group) in both the pre-ischemia and post-ischemia injections. Neutrophils were gated as DAPI$^{-}$CD11B$^{+}$Ly6G$^{+}$.

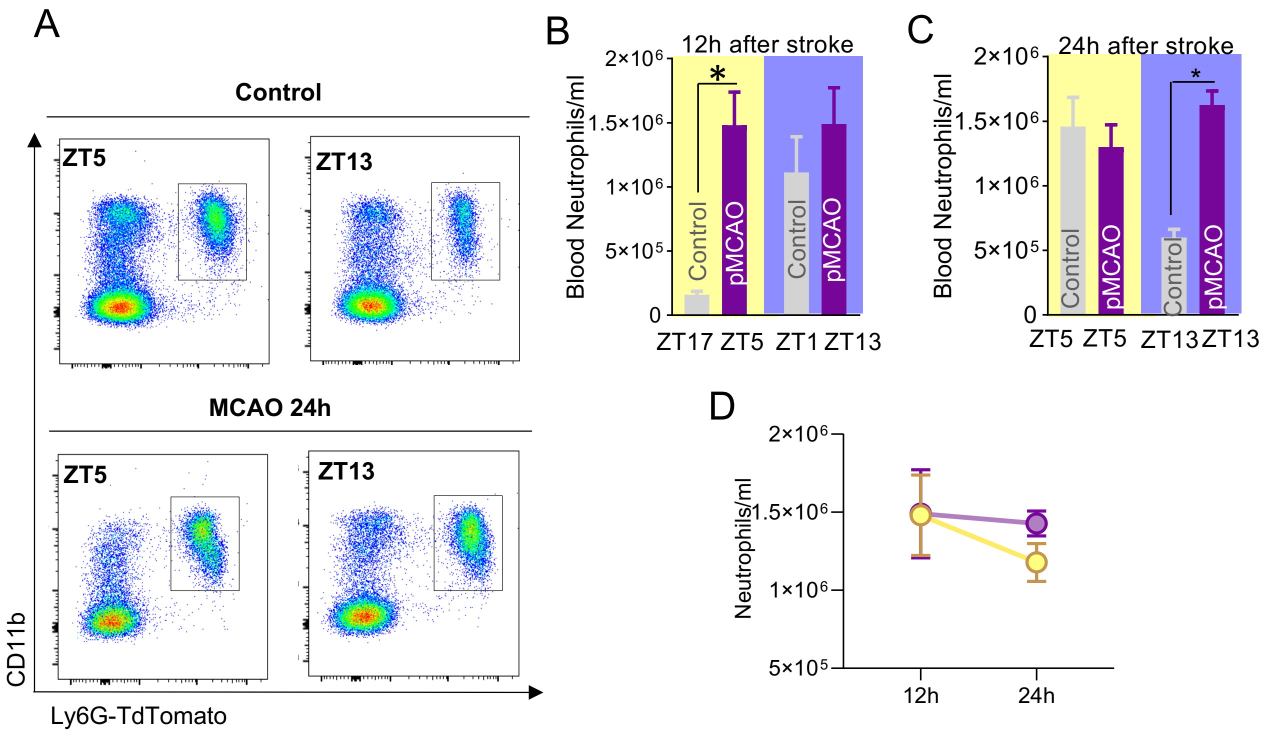


**Supplementary Figure 2. Analysis of blood neutrophils at different times after stroke.** (A) Representative flow cytometry plots of blood neutrophils from control and pMCAO mice 24 hours after stroke at both ZT5 and ZT13. Neutrophils are gated as DAPI-CD45+CD11B+Ly6G+. (B) Blood neutrophil quantification (neutrophils/ml) in both naïve control group (at ZT17 and ZT1 respectively) and 12 hours after MCAO group (subjected to surgery at ZT5 and ZT13). A two-way ANOVA test was performed, considering a p-value < 0.05 as significant * indicates comparison vs. control. (C) Blood neutrophil quantification (neutrophils/ml) in both naïve control group (at ZT5 and ZT13 respectively) and 24 hours after MCAO group (stroke at ZT5 and ZT13 respectively). A two-way ANOVA test was performed, considering a p-value < 0.05 as significant * indicates comparison vs. control. (D) Comparison of blood neutrophil numbers 12 and 24h after stroke in mice subjected to surgery at ZT5 (in yellow) and ZT13 (in purple).

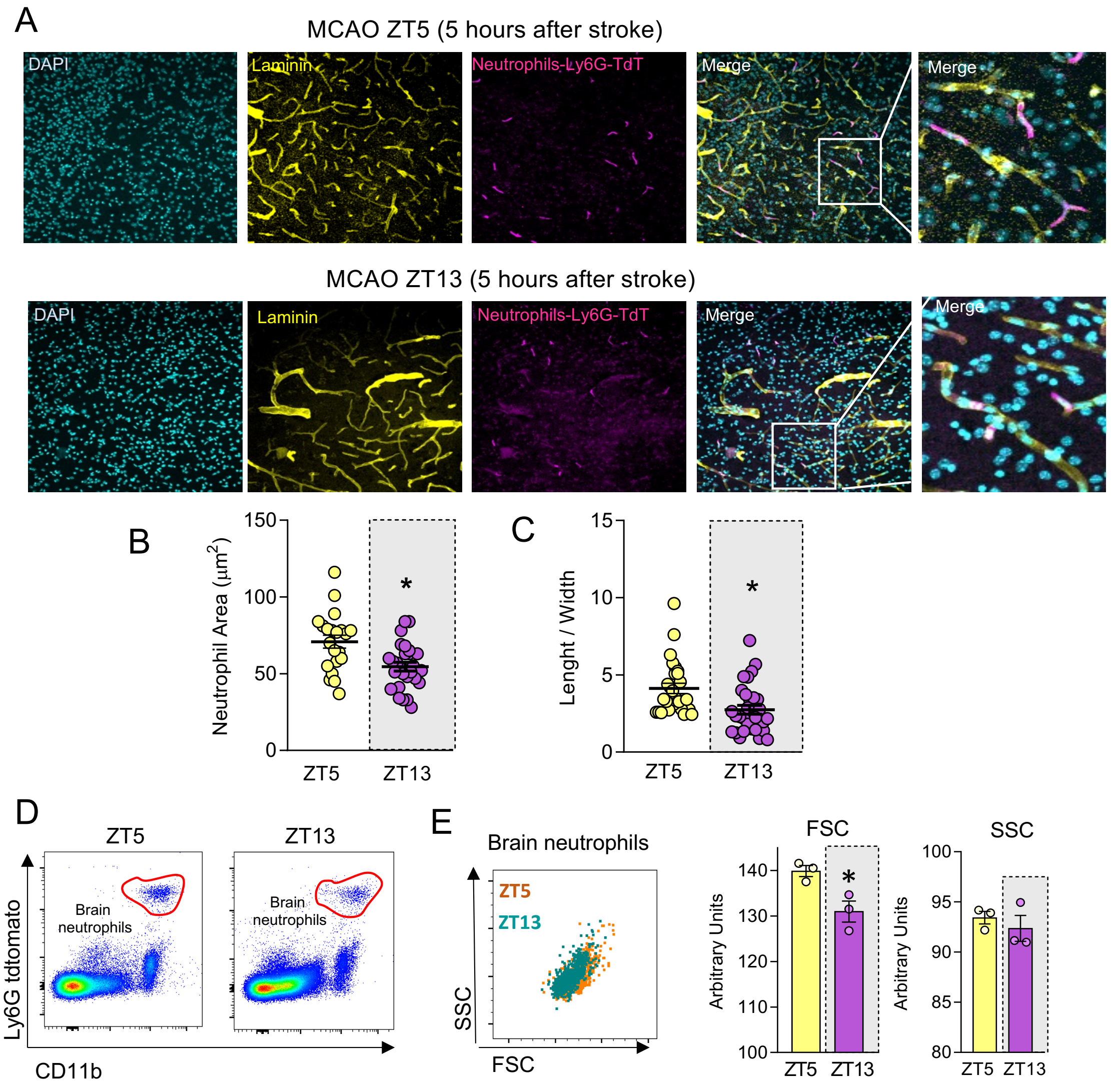


**Supplementary Figure 3. Analysis of brain neutrophils 5 hours after stroke.** (A) Representative immunofluorescence images of brain neutrophils 5 hours post-MCAO, taken from both ZT5 and ZT13 ischemic mice. Nuclei stained with DAPI and are shown in cyan, blood vessels labeled with laminin are shown in yellow, and neutrophils expressing TdTomato from Catchup mice are shown in magenta. (B) Quantification of the size of brain neutrophils estimated as the area (µm$^2$) 5 hours post-MCAO, taken at both ZT5 and ZT13. A non-parametric test was performed, considering a p-value $< 0.05$ as significant. * vs. ZT5. (C) Estimation of the length / width ratio for each brain neutrophil from ZT5 and ZT13 ischemic mice 5h after stroke. A non-parametric test was performed, considering a p-value $< 0.05$ as significant. * vs. ZT5. (D) Representative flow cytometry plots of brain neutrophils from ZT5 and ZT13 ischemic mice 5h after stroke. Neutrophils are gated as DAPI$^-$ CD45$^+$CD11B$^+$Ly6G$^+$. (E) Representative flow cytometry plots displaying SSC and FSC parameters of brain neutrophils from ZT5 and ZT13 ischemic mice 5h after stroke. (F-G) Quantification of the mean FSC and SSC parameters estimated as arbitrary units (AU) in brain neutrophils from ZT5 and ZT13 ischemic mice 5 hours post-MCAO. Data, represented as mean ± SEM, were compared by Mann-Whitney t-test for two groups.

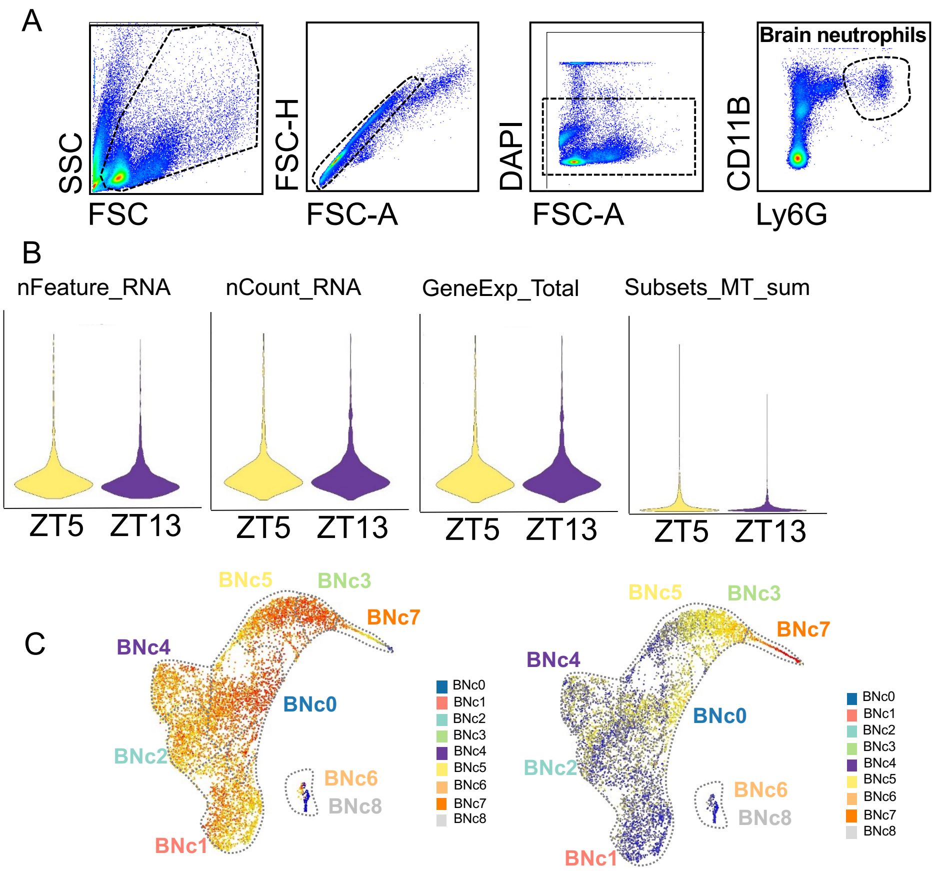


**Supplementary Figure 4. Quality assessment of brain neutrophils for scRNA-seq analysis.** (A) Gating strategy for sorting of brain neutrophils 12h after stroke in both ZT5 and ZT13 ischemic mice. Neutrophils were sorted as $CD45^{hi}$, $CDb11^{+}$, $Ly6G^{+}$, $DAPI^{-}$. (B) Violin plots of the number of genes, feature RNA, RNA counts, number of UMIs and percentage of mitochondrial UMIs. (C) UMAPs of the 3351 brain neutrophils colored by z-score scaled expression showing that BNc7 is the one with high bone marrow proximity score (right) and low maturation score (left).

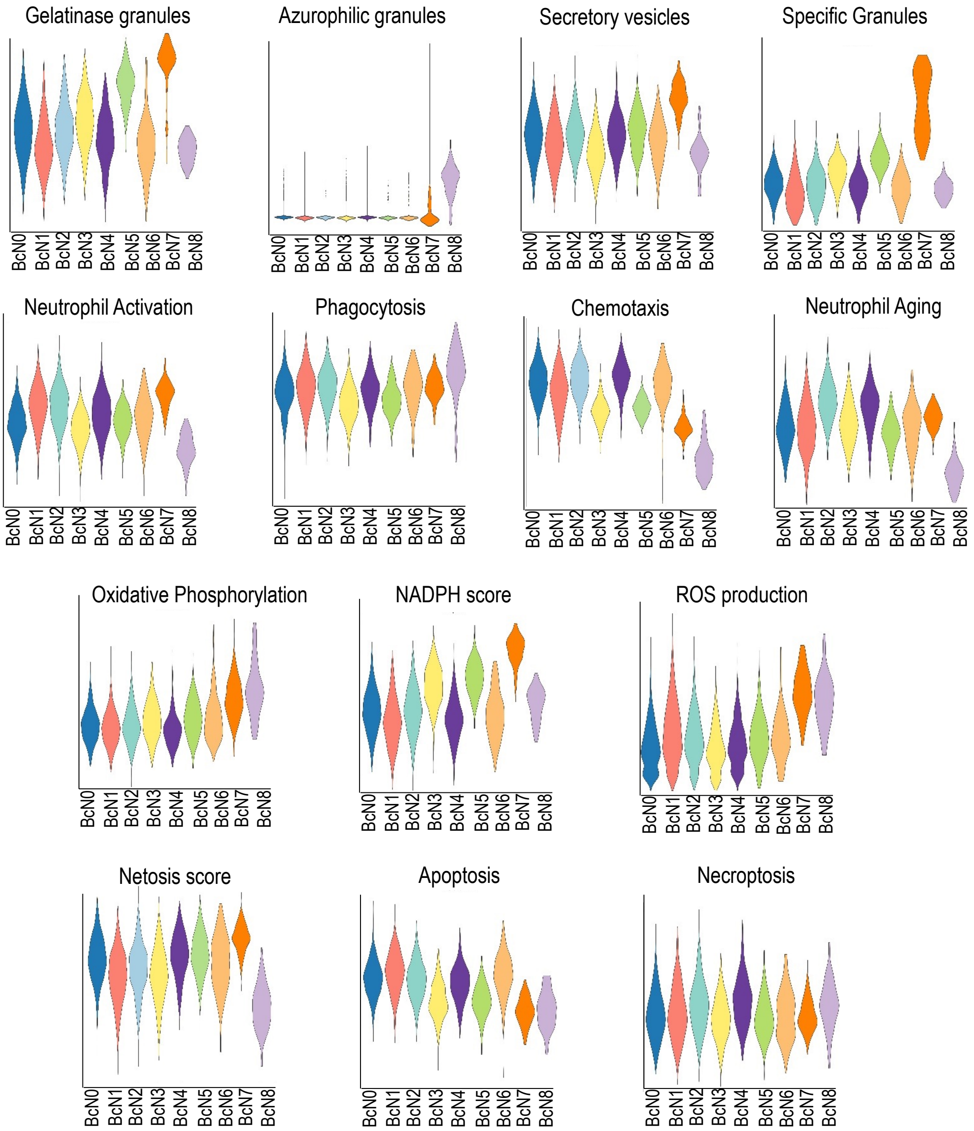


**Supplementary Figure 5. Transcriptional profile of different brain neutrophil subsets for various core and effector neutrophil functions.** Violin plots of different scores for neutrophil effector functions defined as weighted average Z-scores of effector function-related genes previously described in the literature (Extended Data Table S2) for different BNCs detected in the brain.

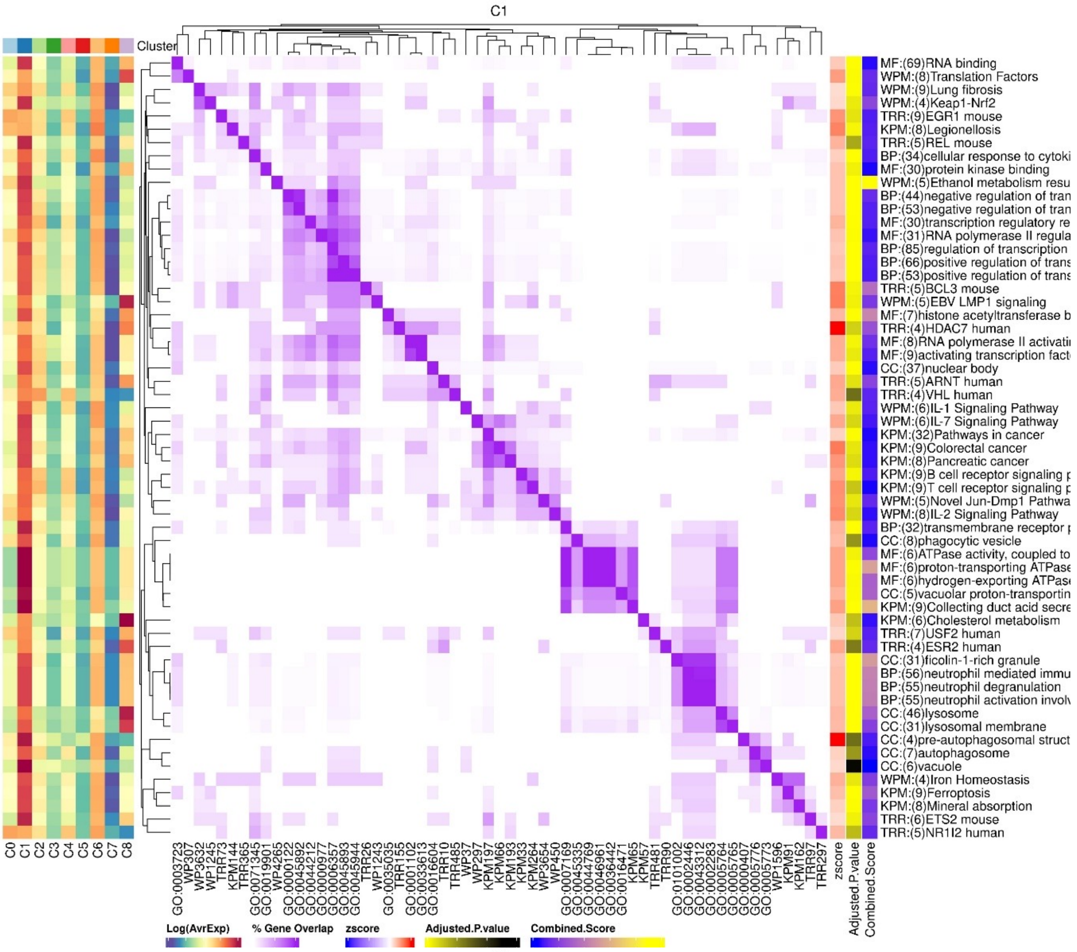
C1
Cluster
MF:(69)RNA binding
WPM:(8)Translation Factors
WPM:(9)Lung fibrosis
WPM:(4)Keap1-Nrf2
TRR:(9)EGR1 mouse
KPM:(8)Legionellosis
TRR:(5)REL mouse
BP:(34)cellular response to cytoki
MF:(30)protein kinase binding
WPM:(5)Ethanol metabolism resu
BP:(44)negative regulation of tran
BP:(53)negative regulation of tran
MF:(30)transcription regulatory re
MF:(31)RNA polymerase II regula
BP:(85)regulation of transcription
BP:(66)positive regulation of trans
BP:(53)positive regulation of trans
TRR:(5)BCL3 mouse
WPM:(5)EBV LMP1 signaling
MF:(7)histone acetyltransferase b
TRR:(4)HDAC7 human
MF:(8)RNA polymerase II activati
MF:(9)activating transcription fact
CC:(37)nuclear body
TRR:(5)ARNT human
TRR:(4)VHL human
WPM:(6)IL-1 Signaling Pathway
WPM:(6)IL-7 Signaling Pathway
KPM:(32)Pathways in cancer
KPM:(9)Colorectal cancer
KPM:(8)Pancreatic cancer
KPM:(9)B cell receptor signaling p
KPM:(9)T cell receptor signaling p
WPM:(5)Novel Jun-Dmp1 Pathwa
WPM:(8)IL-2 Signaling Pathway
BP:(32)transmembrane receptor p
CC:(8)phagocytic vesicle
MF:(6)ATPase activity, coupled to
MF:(6)proton-transporting ATPase
MF:(6)hydrogen-exporting ATPase
CC:(5)vacuolar proton-transportin
KPM:(9)Collecting duct acid secre
KPM:(6)Cholesterol metabolism
TRR:(7)USF2 human
TRR:(4)ESR2 human
CC:(31)ficolin-1-rich granule
BP:(56)neutrophil mediated immu
BP:(55)neutrophil degranulation
BP:(55)neutrophil activation involv
CC:(46)lysosome
CC:(31)lysosomal membrane
CC:(4)pre-autophagosomal struct
CC:(7)autophagosome
CC:(6)vacuole
WPM:(4)Iron Homeostasis
KPM:(9)Ferroptosis
KPM:(8)Mineral absorption
TRR:(6)ETS2 mouse
TRR:(5)NR1I2 human
C0 C1 C2 C3 C4 C5 C6 C7 C8
zscore
Adjusted.P.value
Combined.Score
Log(AvrExp)
% Gene Overlap
zscore
Adjusted.P.value
Combined.Score

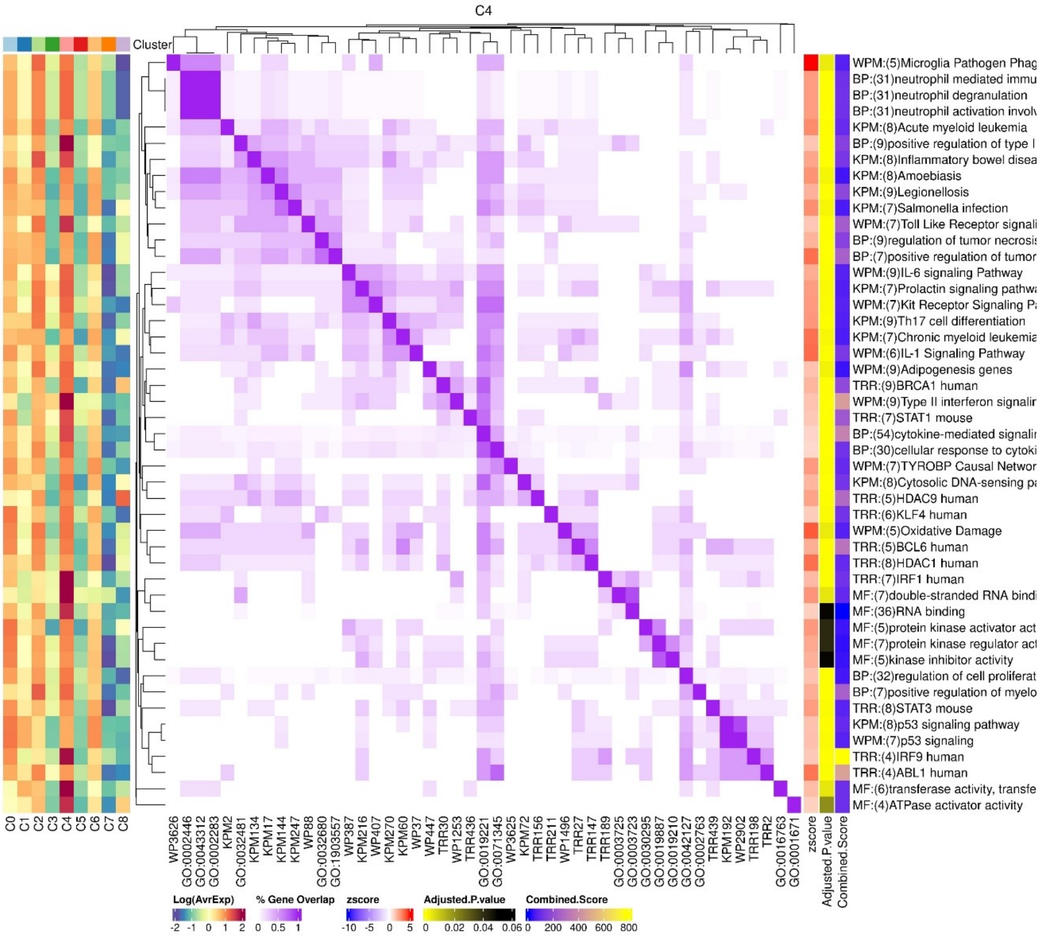
C4
Cluster
WPM:(5)Microglia Pathogen Phag
BP:(31)neutrophil mediated immu
BP:(31)neutrophil degranulation
BP:(31)neutrophil activation involv
KPM:(8)Acute myeloid leukemia
BP:(9)positive regulation of type I
KPM:(8)Inflammatory bowel disea
KPM:(8)Amoebiasis
KPM:(9)Legionellosis
KPM:(7)Salmonella infection
WPM:(7)Toll Like Receptor signali
BP:(9)regulation of tumor necrosis
BP:(7)positive regulation of tumor
WPM:(9)IL-6 signaling Pathway
KPM:(7)Prolactin signaling pathwa
WPM:(7)Kit Receptor Signaling Pa
KPM:(9)Th17 cell differentiation
KPM:(7)Chronic myeloid leukemia
WPM:(6)IL-1 Signaling Pathway
WPM:(9)Adipogenesis genes
TRR:(9)BRCA1 human
WPM:(9)Type II interferon signalin
TRR:(7)STAT1 mouse
BP:(54)cytokine-mediated signalin
BP:(30)cellular response to cytoki
WPM:(7)TYROBP Causal Networ
KPM:(8)Cytosolic DNA-sensing pa
TRR:(5)HDAC9 human
TRR:(6)KLF4 human
WPM:(5)Oxidative Damage
TRR:(5)BCL6 human
TRR:(8)HDAC1 human
TRR:(7)IRF1 human
MF:(7)double-stranded RNA bindi
MF:(36)RNA binding
MF:(5)protein kinase activator act
MF:(7)protein kinase regulator act
MF:(5)kinase inhibitor activity
BP:(32)regulation of cell proliferat
BP:(7)positive regulation of myelo
TRR:(8)STAT3 mouse
KPM:(8)p53 signaling pathway
WPM:(7)p53 signaling
TRR:(4)IRF9 human
TRR:(4)ABL1 human
MF:(6)transferase activity, transfe
MF:(4)ATPase activator activity
C0 C1 C2 C3 C4 C5 C6 C7 C8
zscore
Adjusted.P.value
Combined.Score
Log(AvrExp)
% Gene Overlap
zscore
Adjusted.P.value
Combined.Score

**Supplementary Figure 6. Functional gene ontology analysis of DEGs in clusters BNC1 (top) AND BNC4 (bottom).** Heatmaps hierarchical clustering of gene expression levels and gene ontology analysis showing the correlation matrix of GO analysis for each cluster. The dendrogram shows the Euclidian distance between samples, while the heatmap shows the Pearson correlation (purple = 1, white = −1).

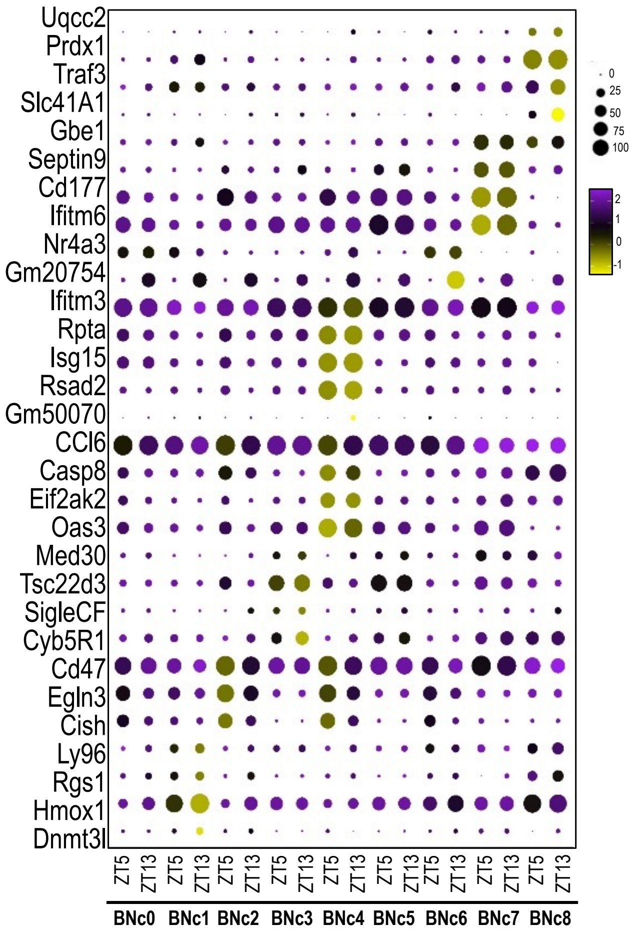


**Supplementary Figure 7. Differentially expressed genes of brain neutrophil ZT5 and ZT13 subsets.** Dot plot showing the scaled expression of signature genes for each cluster that significantly changes between ZT5 and ZT13 ischemic mice, colored by the average expression of each gene in each cluster scaled across all clusters. Dot size represents the percentage of cells in each cluster with more than one read of the corresponding gene.

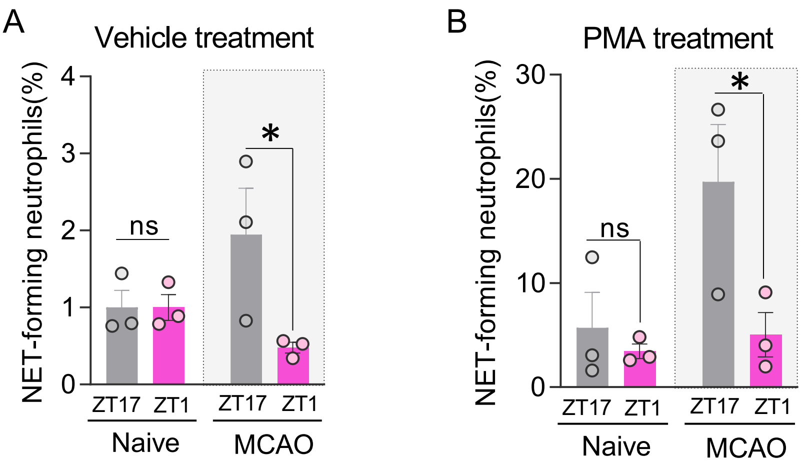


**Supplementary Figure 8. NETosis as a circadian-dependent mechanism promoting microthrombosis after stroke.** (A-B) Quantification of NET-forming neutrophils (%) in vehicle (A) and PMA treatment (B) in both naïve control group (at ZT17 and ZT1) and 12 hours after ZT5 and ZT13 MCAO group in a NETosis *ex vivo* assay. A two-way ANOVA test was performed followed by Bonferroni, considering p<0.05 as significant. *vs. naive.

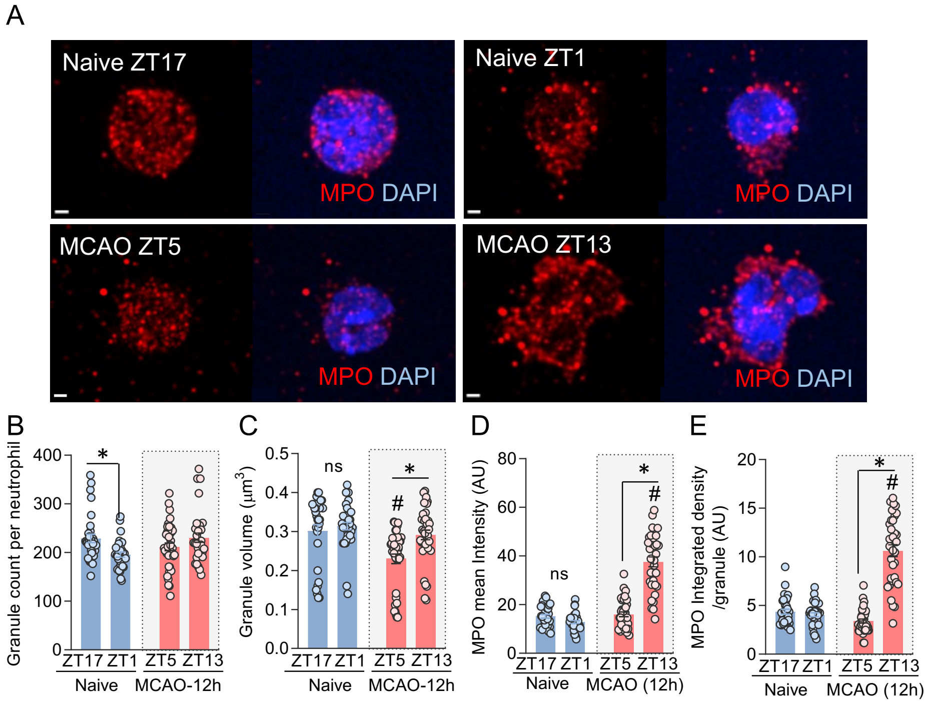


**Supplementary Figure 9. Myeloperoxidase⁺ granule quantification from blood isolated neutrophils after stroke.** (A) Representative immunofluorescence images of blood neutrophils from control group (ZT17 and ZT1 naïve mice) and 12h after MCAO group (stroke at ZT5 and ZT13). MPO is in red and nuclei, stained with DAPI, are in blue. (B) Quantification of the number of MPO granules per neutrophil, (C) the mean granule volume ($\mu m^3$) per neutrophil, (D) the MPO mean intensity measured as arbitrary units (AU), and (E) the MPO integrated density per granule (AU) in both naïve control group (at ZT17 and ZT1) and MCAO group (stroke at ZT5 and ZT13 respectively) analyzed 12h after stroke. A two-way ANOVA test was performed followed by Bonferroni post-hoc. *p<0.05.

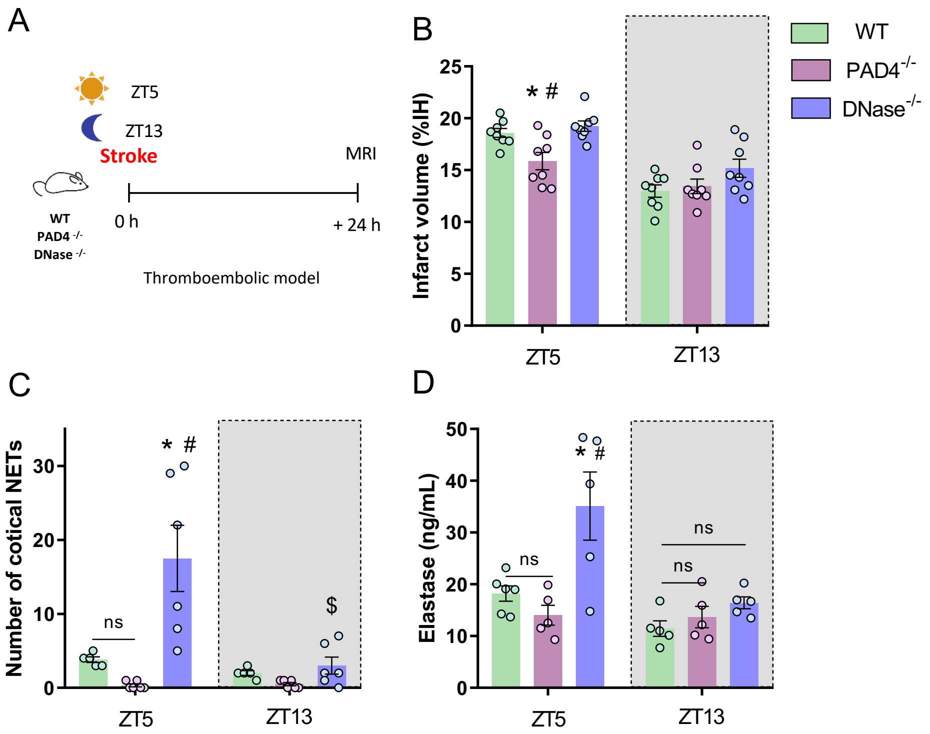


**Supplementary Figure 10. Effect of the modulation of NETosis in an *in situ* thromboembolic model.** (A) Quantification of the percentage of infarcted hemisphere in a *in situ* thromboembolic model at both ZT5 and ZT13 in WT, PAD4$^{-/-}$ and DNase1/3$^{-/-}$ mice. Data were compared by two-way ANOVA test followed by a Bonferroni post-hoc test considering * and # p-value < 0.05 as significant wherein * indicates comparison vs. ZT5 WT and # indicates comparison vs. ZT5 DNase$^{-/-}$. (B) Number of cortical NETs in the *in situ* thromboembolic model both at ZT5 and ZT13 in WT group, PAD4$^{-/-}$ group and DNase$^{-/-}$ group. Data were compared by two-way ANOVA test followed by a Bonferroni post-hoc test considering * and # p-value < 0.05 as significant wherein * indicates comparison vs. ZT5 WT and # indicates comparison vs. ZT5 DNase$^{-/-}$. (C) Elastase quantification (ng/mL) in the thromboembolic model at ZT5 and ZT13 in WT, PAD4$^{-/-}$ and DNase$^{-/-}$ groups. A two-way ANOVA test was performed, considering a p-value < 0.05 as significant. * vs. ZT5 WT; # vs. ZT5 PAD4$^{-/-}$.

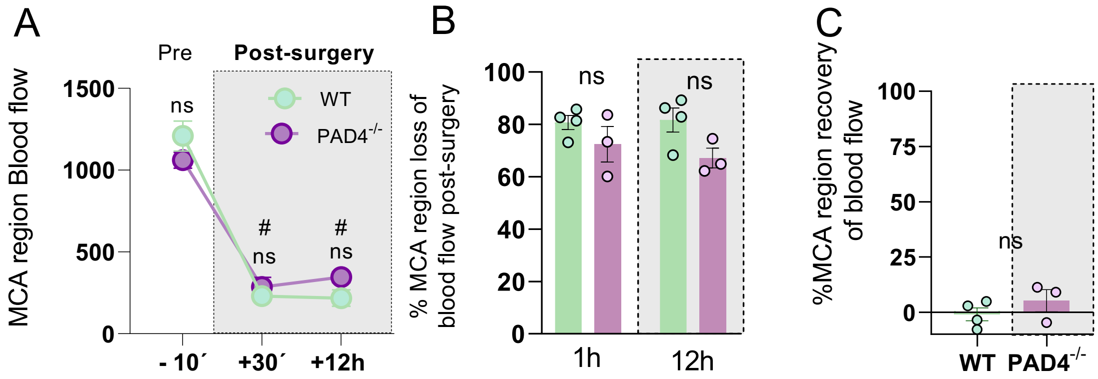


**Supplementary Figure 11. Evaluation of cerebral blood flow in a NETosis deficient model caused by genetic deletion of PAD4.** (A) MCA region blood flow quantification 10 minutes before stroke (-10‘), 30 minutes after stroke (+30‘) and 12 hours after stroke (+12 h) in WT and PAD4$^{-/-}$ animals at ZT5. A two-way ANOVA test was used, and results were considered statistically significant when the p-value was < 0.05 where # indicates statistical significance compared to the pre-surgery condition. (B) Percentage of blood flow loss in MCA region both 30 minutes and 12 hours after ischemia compared to the pre-surgery time point at ZT5 in both WT and PAD4$^{-/-}$ group. A two-way ANOVA test was used, and results were considered statistically significant when the p-value was < 0.05. (C) Percentage of blood flow recovery 12 hours after ischemia (12 h) compared to the pre-surgery time point (-10’) at ZT5 in both WT and PAD4$^{-/-}$ group. A non-parametric test was used, and results were considered statistically significant when the p-value was < 0.05.

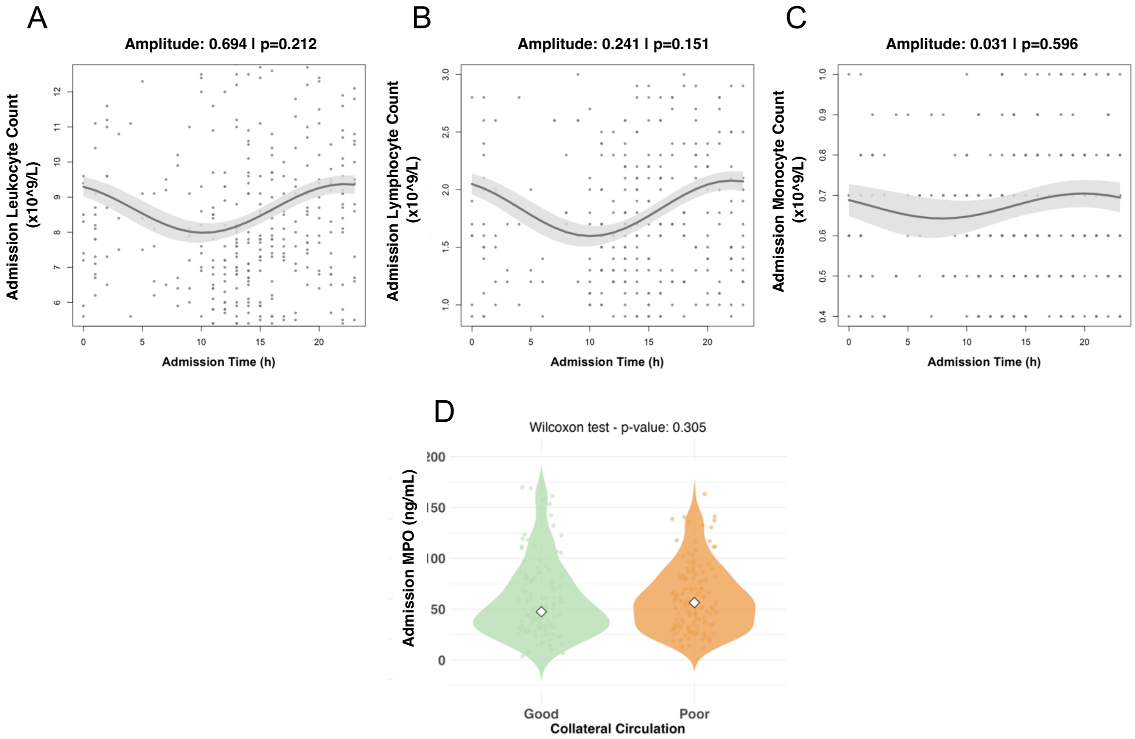


**Supplementary Figure 12. Stroke patients display diurnal oscillations in stroke severity, immunothrombosis and NETosis markers linked to collateral circulation.** (A-C) Diurnal oscillations of admission hematological parameters including leukocytes, lymphocytes and monocytes. Each plot presents the COSINOR-fitted curve with a shaded ribbon representing the standard error (SE) of the fit. The amplitude and p-value of each oscillation are indicated in the plot titles; rhythms with p-values below 0.05 are considered to have significant diurnal patterns. (D) Comparative analysis of clinical markers by collateral circulation status using violin plots. Violin plots illustrate the distribution of MPO in patients with good versus poor collateral circulation (n= 349). Violin plot includes data points (jittered) and displays the median as a white diamond. Outliers were removed using the interquartile range (IQR) method, with outliers defined as values lying beyond 1.5 times the IQR from the first and third quartiles. For each marker, appropriate statistical tests were conducted based on data distribution. t-test applied to admission elastase and Wilcoxon tests applied to admission NIHSS and infarct size.

| FOOTPRINT (parameter) | | ZT5 Baseline | | ZT5 MCAO | | ZT13 Baseline | | ZT13 MCAO | |
|---|---|---|---|---|---|---|---|---|---|
| | | **MEAN** | SD | **MEAN** | SD | **MEAN** | SD | **MEAN** | SD |
| **Hindlimb stride lenght (cm)** | Contralateral (right) | **6.938** | 0.211 | **6.821** | 0.720 | **6.983** | 0.914 | **7.024** | 0.807 |
| | Ipsilateral (left) | **7.056** | 0.328 | **6.893** | 0.792 | **6.905** | 0.638 | **6.996** | 0.803 |
| **Sway lenght (cm)** | Forelimb | **1.641** | 0.235 | **1.862** | 0.138 | **1.607** | 0.173 | **1.577** | 0.147 |
| | Hindlimb | **2.619** | 0.415 | **2.659** | 0.136 | **2.651** | 0.136 | **2.369** | 0.147 |
| **Overlap (average forelimb and hindlimb) cm** | | **0.762** | 0.370 | **0.726** | 0.359 | **0.659** | 0.255 | **0.527** | 0.304 |

**Supplementary Table S1. Evaluation of motor impairment after stroke by footprint test.** Main parameters measured in the footprint test: hindlimb stride length (cm), sway length (cm) and overlap (average forelimb and hindlimb; cm). All these parameters were measured in ZT5 and ZT13 mice during the baseline (48h before stroke) and 24h after stroke.

**Supplementary Table S2. Enriched pathways in differentially expressed genes from each cluster**

**Supplementary Table S3. Differentially expressed genes in WT Brain at ZT13 vs. ZT5 per cluster**

**Supplementary Table S4. Enriched pathways in differentially expressed genes in WT Brain at ZT13 vs. ZT5 per cluster**

| n | Overall<br>377 |
|---|---|
| Age, mean (SD) | 76.00 [63.50, 84.00] |
| Gender (Female), n (%) | 188 (51.2) |
| Previous mRS, n (%) | |
| mRS 0 | 283 (77.1) |
| mRS 1 | 51 (13.9) |
| mRS 2 | 33 (9.0) |
| Tobacco, n (%) | |
| Non smoker | 247 (68.4) |
| Active smoker | 64 (17.7) |
| Past smoker | 50 (13.9) |
| Alcohol, n (%) | |
| Non Alcohol Use | 311 (87.1) |
| Alcohol Use | 34 (9.5) |
| Past Alcohol Use | 12 (3.4) |
| Hypertension, n (%) | 250 (66.3) |
| Dyslipidemia, n (%) | 216 (57.3) |
| Diabetes, n (%) | 95 (25.9) |
| Atrial Fibrillation, n (%) | 97 (25.7) |
| Metallic Valve, n (%) | 15 (4.1) |
| Previous Ischemic Stroke, n (%) | 55 (14.6) |
| Previous Hematoma, n (%) | 5 (1.3) |
| Previous Myocardial Infarction, n (%) | 37 (10.1) |
| Arteriopathy, n (%) | 27 (7.2) |
| Chronic Kidney Disease, n (%) | 22 (6.0) |
| Previous Antihypertensive Therapy, n (%) | 230 (61.0) |
| Previous Lipid-Lowering Therapy, n (%) | 189 (50.1) |
| Previous Antiplatelet Therapy, n (%) | 88 (23.3) |
| Previous Anticoagulant Therapy, n (%) | 85 (22.5) |

**Supplementary Table 5.** Baseline characteristics of ischemic stroke cohort with known onset (n=377). Results are presented as median [IQR] or count (%).

| n | Overall<br>377 |
|---|---|
| Stroke Onset-to-Hospital Time, mean (SD) | 2.00 [1.00, 4.00] |
| Admission Blood Glucose, mean (SD) | 118.00 [103.00, 143.00] |
| Admission Leukocytes, mean (SD) | 8.10 [6.80, 9.90] |
| Admission Neutrophils, mean (SD) | 5.50 [4.10, 7.00] |
| Admission Lymphocytes, mean (SD) | 1.70 [1.20, 2.30] |
| Admission Monocytes, mean (SD) | 0.60 [0.50, 0.80] |
| Admission Platelets, mean (SD) | 211.00 [170.25, 260.75] |
| Admission MPV, mean (SD) | 9.00 [8.30, 9.70] |
| Admission Neutrophil-Specific Elastase, mean (SD) | 37.93 [27.11, 56.71] |
| Admission Myeloperoxidase, mean (SD) | 57.30 [36.12, 89.21] |
| Admission sCD40L, mean (SD) | 0.30 [0.15, 0.55] |
| Admission NIHSS, mean (SD) | 10.00 [4.00, 18.00] |
| Infarct Volume (cc), mean (SD) | 3.58 [0.25, 18.88] |
| Collateral Circulation, mean (SD) | |
| Poor | 48 (21.1) |
| Intermediate | 81 (35.5) |
| Good | 99 (43.4) |
| Etiology | |
| Atherothrombotic | 116 (31.8) |
| Cardioembolic | 142 (38.9) |
| Cryptogenic | 28 (7.7) |
| Coexistence of Cause | 31 (8.5) |
| Incomplete Study | 28 (7.7) |
| Unusual Cause | 20 (5.5) |
| tPA, n (%) | 163 (45.8) |
| Thrombectomy, n (%) | 215 (60.1) |
| Post-hemorrhage, n (%) | 57 (16.8) |
| NIHSS at Stroke Unit, mean (SD) | 3.00 [1.00, 9.00] |
| mRS 3 months | |
| Independent | 219 (63.7) |
| Dependent | 83 (24.1) |
| All-cause mortality | 42 (12.2) |

**Supplementary Table 6.** Clinical characteristics, therapeutic management, and prognostic indicators of the ischemic stroke cohort (n=377). Results are presented as median [IQR] or count (%).

| Biomarker | MESOR ± SE | Amplitude ± SE | Amplitude p-value | Acrophase (horas) ± SE | Acrophase p-value | Peak max (value) | Peak max (h) |
|---|---|---|---|---|---|---|---|
| ***Cosinor model fitted based on time of admission*** | | | | | | | |
| **Admission Leukocytes** | 8.678 ± 0.167 | 0.694 ± 0.204 | **0.001** | 22.189 ± 1.360 | 0.183 | 9,296 | 22 |
| **Admission Neutrophils** | 5.984 ± 0.163 | 0.402 ± 0.199 | **0.043** | 22.281 ± 2.299 | 0.455 | 6,346 | 22 |
| **Admission Lymphocytes** | 1.838 ± 0.053 | 0.241 ± 0.066 | **0.000** | 22.067 ± 1.249 | 0.122 | 2,049 | 22 |
| **Admission Platelets** | 218.786 ± 4.340 | 15.681 ± 5.573 | **0.005** | 21.414 ± 1.509 | 0.087 | 231,011 | 21 |
| **Admission Mean Platelet Volume** | 9.047 ± 0.065 | 0.178 ± 0.078 | **0.022** | 12.691 ± 2.063 | 0.738 | 9,222 | 13 |
| **Admission sCD40L** | 0.427 ± 0.028 | 0.089 ± 0.036 | **0.014** | 21.086 ± 1.650 | 0.088 | 0,493 | 21 |
| **Admission log Neutrophil-Elastase** | 3.678 ± 0.036 | 0.096 ± 0.055 | 0.079 | 17.619 ± 1.692 | **0.001** | 3,688 | 18 |
| **Admission log Myeloperoxidase** | 3.995 ± 0.048 | 0.200 ± 0.071 | **0.005** | 18.977 ± 1.104 | **0.000** | 4,046 | 19 |
| ***Cosinor model fitted based on time of symptom onset*** | | | | | | | |
| **Admission NIHSS** | 11.480 ± 0.493 | 1.534 ± 0.591 | **0.009** | 20.020 ± 1.841 | **0.031** | 12,253 | 20 |
| **24h post-stroke NIHSS** | 5.946 ± 0.432 | 1.150 ± 0.619 | 0.063 | 5.649 ± 1.830 | **0.002** | 6,051 | 6 |
| **Delta NIHSS (24-admission)** | 42.356 ± 4.454 | 9.391 ± 6.517 | 0.150 | 13.547 ± 2.235 | 0.489 | 50,988 | 14 |
| **Infarct Volume (cc)** | 24.294 ± 3.887 | 12.928 ± 5.957 | **0.030** | 16.358 ± 1.345 | **0.001** | 29,674 | 16 |

**Supplementary Table 7. Table showing diurnal patterns of blood markers at admission and clinical variables in acute stroke patients after COSINOR adjustment.** The table shows the diurnally adjusted estimates (Estimate ± Standard Error) of blood markers at admission and clinical variables in acute stroke patients. Each variable includes MESOR, amplitude, acrophase (in hours), and the corresponding p-value, providing insights into the timing and intensity of diurnal patterns.
